# CDK9 degradation Inhibits Gastroesophageal Cancer growth and Overcomes Radiation Resistance by Increasing Chromatin Accessibility and Downregulating YAP1/TEAD signaling

**DOI:** 10.1101/2025.02.22.639565

**Authors:** Yan-ting Yann Zhang, Mikel Ghelfi, Yu Xue, Haiying Chen, Gennaro Calendo, Generosa Grana, Yuan Li, Dipti Athavale, Curt Balch, Xiaodan Yao, Woonbok Chung, Zhenning Wang, Francis Spitz, Tracy Tang, Vladimir Khazak, Jia Zhou, Jean-Pierre Issa, Shumei Song

**Author notes:** Corresponding author: Yan-ting Yann Zhang, Ph.D; and Shumei Song, MD, PhD.

## Abstract

**Background:** Gastroesophageal cancer (GEAC) remains a major health burden and urgently needs novel therapeutic targets. The inhibition of CDK9’s activity holds the potential to be a highly effective anti-cancer therapeutic. However, the functional role of CDK9, and its potential targeting in GEAC, remain largely unknown.

**Objective:** We aim to evaluate the potential of degradation CDK9 in GEAC treatment and explore its mechanisms.

**Design:** We evaluated the expression and distribution of CDK9 in GEAC tissue. We designed and synthesized novel CDK9 degraders using proteolysis targeting chimeras (PROTACs) strategy and selected the promising one for further anti-tumor activity evaluation both in vitro and in vivo. We evaluated the effects of CDK9 degradation on epigenetic reactivation, gene expression and chromatin accessibility. We evaluated the co-targeting of CDK9 and YAP/Tead signaling for GEAC treatment, especially for radiation-resistant tumor treatment.

**Results:** We demonstrated significantly elevated CDK9 expression in primary GEAC tumor tissues compared to normal tissues, in association with poor survival. We developed a novel CDK9 degrader, YX0597, reducing RNA Pol II Serine 2 phosphorylation, and inhibition MCL-1; this was accompanied by potent inhibition of GEAC cell growth, especially in radiation-resistant tumor cells. Mechanistically, YX0597 strongly enhanced chromatin accessibility, activating epigenetically silenced genes; and dramatically inhibited YAP/TEAD signaling. CDK9 closely interacts with YAP/TEAD signaling, and co-targeting these two mediators could be a novel treatment strategy for the treatment of GEAC.

**Conclusion:** Our studies reveal a new avenue for targeting CDK9-hyperactivated GEAC tumors, especially in combination with YAP1/TEAD inhibition in radiation-resistant GEAC tumors.

**What is already known on this subject?:** Gastroesophageal cancer (GEAC), the third-leading cause of global cancer death and urgently needs novel targeted therapies. Targeted protein degradation has emerged as an attractive strategy to fight cancer, complementing the activity of traditional small-molecule inhibitors. Cyclin-dependent kinase 9 (CDK9) has been implicated in various cancers. Recently, CDK9 degraders have now been developed for future clinical benefit in targeting tumors with highly activated CDK9.

**What are the new findings?:** In this study, we show that CDK9 expression was significantly elevated in primary GEAC tumor tissues including PDXs compared to normal tissues, in association with poor survival. We developed a novel CDK9 degrader XY0597 and proved that it inhibits the progression of GEAC by increasing chromatin accessibility and down-regulating YAP/Tead signaling. We also proved a strong connection between CDK9 and YAP/TEAD signaling and Co-targeting them synergistically inhibits GEAC tumor cell growth, especially in tumors with high YAP signaling activity, such as radiation-resistant cancer.

**How might it impact on clinical practice in the foreseeable future?:** Our data suggests that CDK9 is a viable anticancer target in GEAC. We demonstrate that a novel CDK9 degrader, YX0597, has high potential clinical application for GEAC treatment, especially in radiation-resistant cancer. We reveal the crosstalk of CDK9 and YAP/TEAD signaling, and that co-targeting these two mediators could be a new avenue for targeting CDK9-hyperactivated GEAC primary and radiation-resistant tumors.

## INTRODUCTION

Gastroesophageal cancer (GEAC), encompassing cancers of the stomach and esophagus, represents the fifth-most common malignancy, worldwide. Gastric cancer alone accounts for over 769,000 deaths annually, globally representing the third-most lethal cancer [1]. In Western nations, adenocarcinoma is the most common histopathological subtype, with gastric cancer risk factors including obesity, consumption of smoked foods, and *Helicobacter Pylori* infection [2], while esophageal cancer risks include smoking, alcohol consumption, high sodium intake, gastroesophageal reflux disease (GERD), and Barrett’s esophagus [3].

Cyclin-dependent kinases (CDKs), a family of serine/threonine protein kinases, play a crucial role in regulating the cell cycle, transcription, and various other physiological processes. Specifically, CDKs form complexes with regulatory proteins called cyclins, which are expressed at specific phases of the cell cycle. These cyclin/CDK complexes phosphorylate target proteins, triggering essential events like DNA replication, chromosome segregation, and cell division. Importantly, CDK7, CDK8, CDK9, CDK12, and CDK13 are key transcriptional regulators, associated with transcription factors and RNA polymerase II (RNAPII), with the best-known of these being CDK9. CDK9 dysregulation has been shown to contribute to development of a variety of malignancies such as pancreatic, prostate, and breast cancers [4–6]. However, the role of CDK9 in GEAC, the common and deadly upper GI tumors, is little understood.

Due to its implication in multiple facets of oncogenesis, CDK9 inhibition holds potential as an effective anti-cancer strategy, with considerable effort currently being invested in developing CDK9 inhibitors (CDKIs). These include flavopiridol and roscovitine, first-generation pan-CDKIs [5], CDKI-73, a highly potent small-molecule CDK9 inhibitor used against colorectal cancer [7], and AZD4573, in clinical trials for patients with relapsed or refractory hematological malignancies [8]. Unfortunately, while most of these inhibitors showed promise in preclinical trials, many produced severe side effects or were not effective in patients [5].

Previously, we reported that CDK9 inhibition reactivated tumor suppressor gene (TSG) expression, via epigenetic regulation, including heterochromatin recruitment of BRG1 [9], a nucleosome remodeler that facilitates gene transcription, derepressing TSGs previously silenced by promoter hypermethylation [10]. Consistent with our previous report, recent research reported that CDK9 inhibition induces epigenetic reprogramming, revealing strategies to circumvent lymphoma resistance [11], suggesting a role for CDK9 in tumorigenic chromatin condensation. CDK9 inhibition can also rapidly alter the transcriptome and proteome, via decreased phosphorylation of Serine 2 in the carboxyl-terminal domain of RNAPII. This event results in RNAPII pausing and suppression of multiple oncogenes (e.g., MYC target genes, HOTAIR lncRNA, PPP1R13L, RBPJ target genes, MCL-1, BIRC5, JunB, PIM3, etc.), via terminated transcriptional elongation [12]. These genes are critical to the survival and proliferation of cancer cells, and it is worth noticing that some of them belonged to the downstream effectors of oncogenic signaling by YAP/TEAD, a negative regulator of the well-known Hippo organ size control pathway (12). However, to date, the association of CDK9 expression with YAP/TEAD signaling has not been explored in GEAC.

More recently, targeted protein degradation has emerged as an attractive strategy to fight cancer, with proteolysis-targeting chimeras (PROTACs, or protein degraders) representing tools for the selective degradation of cancer targets (including CDK9), complementing the activity of traditional small-molecule inhibitors [13]. These compounds typically incorporate previously reported inhibitors, and a known E3 ligase ligand, to induce ubiquitination, and subsequent degradation, of a target protein [13]. CDK9 degraders have now been developed for future clinical benefit in targeting tumors with highly activated CDK9 [14].

In this study, we developed a novel CDK9 degrader XY0597 and proved that it inhibits the progression of GEAC by increasing chromatin accessibility and down-regulating YAP/Tead signaling. We also proved a strong connection between CDK9 and YAP/TEAD signaling and Co-targeting them synergistically inhibits GEAC tumor cell growth, especially for irradiation resistant tumor. Taken together, our studies have opened a new avenue for targeting CDK9 hyperactivated GEAC tumors especially in combination with YAP1/TEAD inhibition.

## RESULTS

### CDK9 is highly expressed in tumor tissues of human primary GEAC and of PDXs

CDK9 is a critical kinase that regulates transcription of several anti-apoptotic and oncogenes, essential for cancer maintenance, growth, metastasis and chemoresistance, whose overactivity has been shown in numerous malignancies such as pancreatic, prostate, and breast cancers [12]. To determine if CDK9 expression is elevated in GEAC and potentially be tumor target, first we explore The Cancer Genome Atlas (TCGA) dataset and found that CDK9 expression was significantly elevated in both esophageal adenocarcinoma (EAC) and gastric adenocarcinoma (GAC), compared to adjacent normal tissues (https://cistrome.shinyapps.io/timer/). (Fig.1A and Fig, S1A). Moreover, *CDK9* gene amplification represented the most common genetic alteration in both EAC and GAC (Fig.S1B), and high CDK9 expression significantly associated with poor survival in GAC (Fig. 1B). Furthermore, we examined CDK9 expression by immunohistochemistry (IHC) staining in GAC TMA containing 421 cases of tumor tissues and 408 cases of normal adjacent tissues (NATs) and showed that CDK9 expression was significantly increased in tumor tissues 81.0% (340/421) compared to NATs 51.0% (208/408), (*P<0.001*; Cut-off value is 1.5) in both intestinal/diffuse type human GAC (Fig. 1C). We further validated the expression of CDK9 in 9 GAC PDX tissues and nuclear CDK9 staining were shown in three representative PDXs (Fig. S2A). Altogether, these data support highly expressed CDK9 could be a therapeutic target in GEAC.

**Figure 1.**
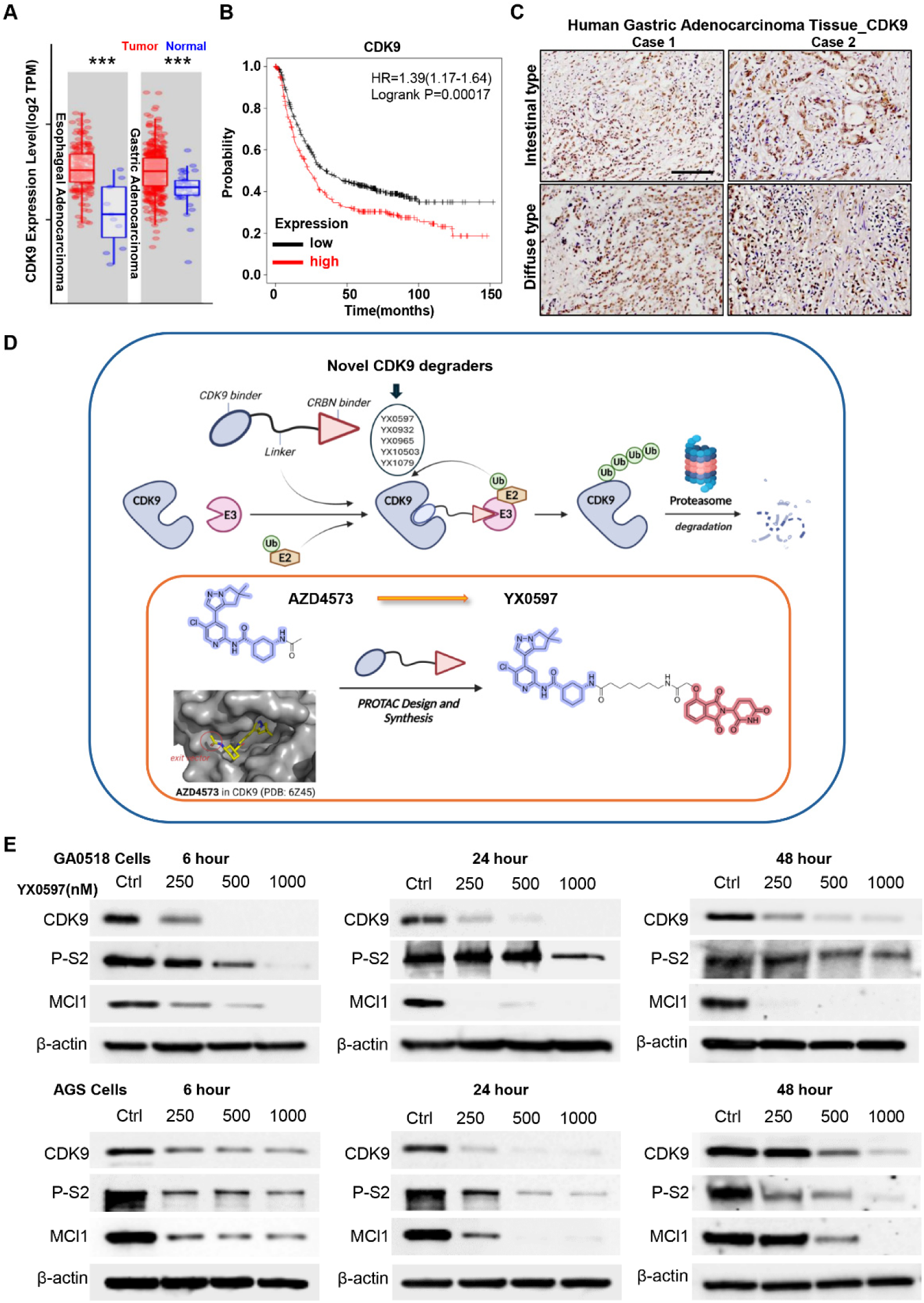
CDK9 is overexpressed in GEAC cancers and is degraded by a novel degrader YX0597. Public datasets showing: **A,** CDK9 expression in gastric (GEAC) and esophageal (EAC) adenocarcinomas versus normal tissues; **B,** Survival analysis of CDK9^high^ vs. CDK9^low^ gastric adenocarcinoma patients (https://kmplot.com/). **C,** Immunohistochemistry of CDK9 expression in intestinal and diffuse human gastric adenocarcinomas. Scale bar:100µm. **D,** Schema of design of novel CDK9-degraders. A CDK9-binding molecule recruits an E3 ligase, via a CRBN binder, resulting in CDK9 ubiquitination and proteasomal degradation. For novel CDK9 degrader YX0597, we chose a commercial CDK9 inhibitor AZD4573 as the CDK9 binder. **E,** The specific CDK9 degrader YX0597 potently and time-dependently suppresses the expression of CDK9, phosphorylation of Pol II S2, and the anti-apoptotic protein MCL1.

### Develop a novel degrader for targeting CDK9 in GEAC cells

Recently, much interest and effort has been invested in targeting CDK9 for cancer research, with several CDK9 inhibitors showing initial promise in preclinical studies, but limited efficacy in clinical trials. In contrast to small-molecule inhibitors alone, however, targeted protein degradation has emerged as a novel cancer therapeutic. Therefore, we synthesized five CDK9 degraders, based on well-known CDK9 inhibitory molecules (Fig. 1D), and compared their cell proliferation inhibition effects in the gastric cancer cell line AGS and GAC patient derived GA0518 cells [15]. Among the five degraders tested, the degrader, YX0597 exerted the more potent inhibition growth in both AGS and GA0518 tumor cells in a dose and time dependent manner (Fig. S3) by Incucyte real time analysis. For novel CDK9 degrader YX0597, we chose a commercial CDK9 inhibitor AZD4573 as the CDK9 binder(Fig. 1D). In the co-crystal of AZD4573 in CDK9, the acetyl group of AZD4573 in the solvent region of CDK9 can provide an exit vector for the construction of CDK9 PROTACs. For the E3 binder, we chose widely-used CRBN binder, such as thalidomide and its derivatives. For the linker, we chose both flexible and rigid linkers. Other degraders showed either less potent or lack of the dose dependent in both tumor cell lines at the different cell density seeding (2K or 4K cells/well) (Fig.S4). Thus, we focus on XY0597 degrader in the following experiments. In GA0518 and AGS cells, YX0597 dose-dependently induced rapid and stable CDK9 degradation, in parallel with decreased RNA polymerase II S2 phosphorylation and expression of the CDK9 downstream effector MCl1 (Fig. 1F). Western blotting showed that YX0597 similarly inhibited CDK9 signaling in PDX cells (Fig. S2B). Immunofluorescence analysis further showed that CDK9 was markedly downregulated by YX0597 (Fig. S2C). Consistent with MCL1 inhibition, cleaved caspase 3-positive cells were greatly increased in YX0597-treated PDXs (Fig. S2D). These analyses show that XY0597 is a novel and efficacious CDK9 degrader, meriting further evaluation of its anti-tumor activity.

### YX0597 exhibits potent antitumor effects in vitro

Since CDK9 is implicated in numerous events in tumor progression, we examined YX0597 effects on specific tumor properties. In GAC patient ascites-derived GA0518 cells, and the AGS cell line GAC, YX0597 exerted dose-dependent antiproliferative effects (Figs. 2A, 2B, and S4). Similarly, YX0597 potently suppressed esophageal cancer cell proliferation (Fig. S5). Transwell migration assays showed strongly inhibited cell migration (Fig. 2C). Moreover, tumor sphere generation (Fig. 2D) and colony formation (Fig. 2E) were greatly inhibited in YX0597-treated GA0518 and AGS cells. These in vitro results suggest that YX0597 potently inhibits tumor proliferation, metastasis, and targets cancer stem cells.

**Figure 2.**
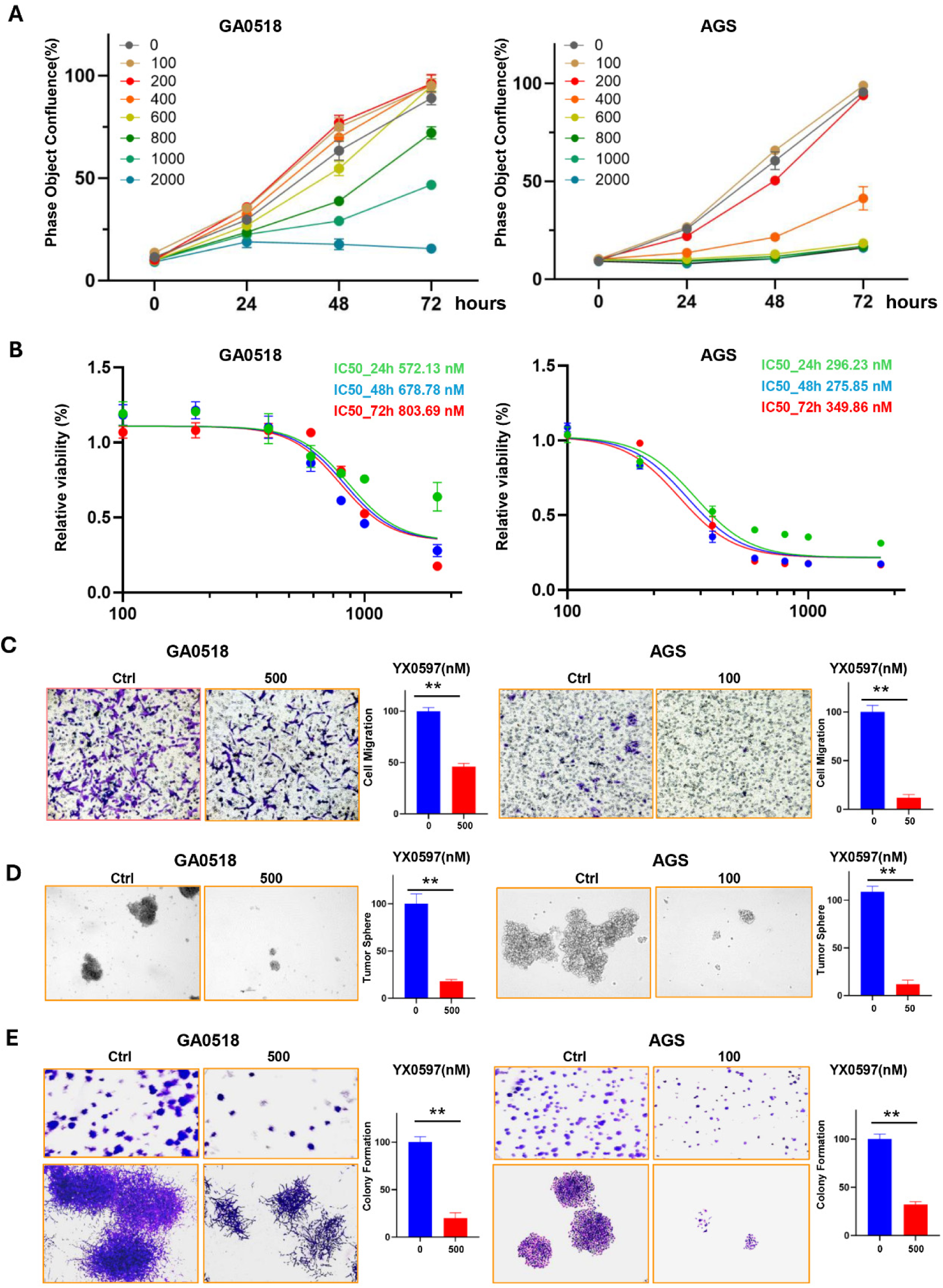
The novel CDK9 degrader XY0597 exerts potent effects in suppressing tumor cell malignant behaviors *in vitro*. **A,** Incucyte live-cell analysis showing dose- and time-dependent growth inhibition by YX0597, in GA0518 and AGS gastric cancer cells. **B,** XY0597 dose-response curve and IC50 values in GA0518 and AGS cells. **C,** Cell migration inhibition by XY0597 treatment for 48 hours. **D,** Inhibition of tumor sphere formation by XY0597 treatment. **E,** Inhibition of colony formation by XY0597. ** *p < 0.01*

### YX0597 epigenetically induces gene expression by increasing chromatin accessibility

We previously used a reporter system, YB5, to show that CDK9 inhibition can reactivate an epigenetically silenced *GFP* gene [9].Here, we likewise used YB5 to test whether a CDK9 degrader can also derepress *GFP*, following CDK9 degradation. Consistently, GFP fluorescence was robustly detected, in a dose-dependent manner, following treatment with YX0597 or YX0965 (Figs. 3A and S6). Thus, either the inhibition of CDK9 activity, or inducing its degradation, could reactivate silenced genes analogously. YX0597 treatment of GA0518 cells, for 24 hours, upregulated over 1000 genes (Fig. 3B). These included 27 tumor-suppressors and gastric cancer biomarkers (Fig. 3C), including *WWOX*, *PINX1*, *IQGAP2*, *MTAP*, and nearly 50 transposable elements (Fig. S7). The gene functional enrichment analysis showed that RNA processing, mitochondria and apoptosis regulation related genes were hugely upregulated (Fig. 3D). Interestingly, plenty of genes functional in cytoskeleton regulation were downregulated, which may contribute to the cell migration inhibition activity of YX0597 (Fig. 3D and Fig.2C). In support of the ability to reactivate silenced genes by YX0597, ATAC-seq analysis showed increased genome-wide chromatin accessibility after YX0597 treatment (Fig. 3D). These data indicate that YX0597 increases gene expression, at the epigenetic level, via increasing chromatin accessibility.

**Figure 3.**
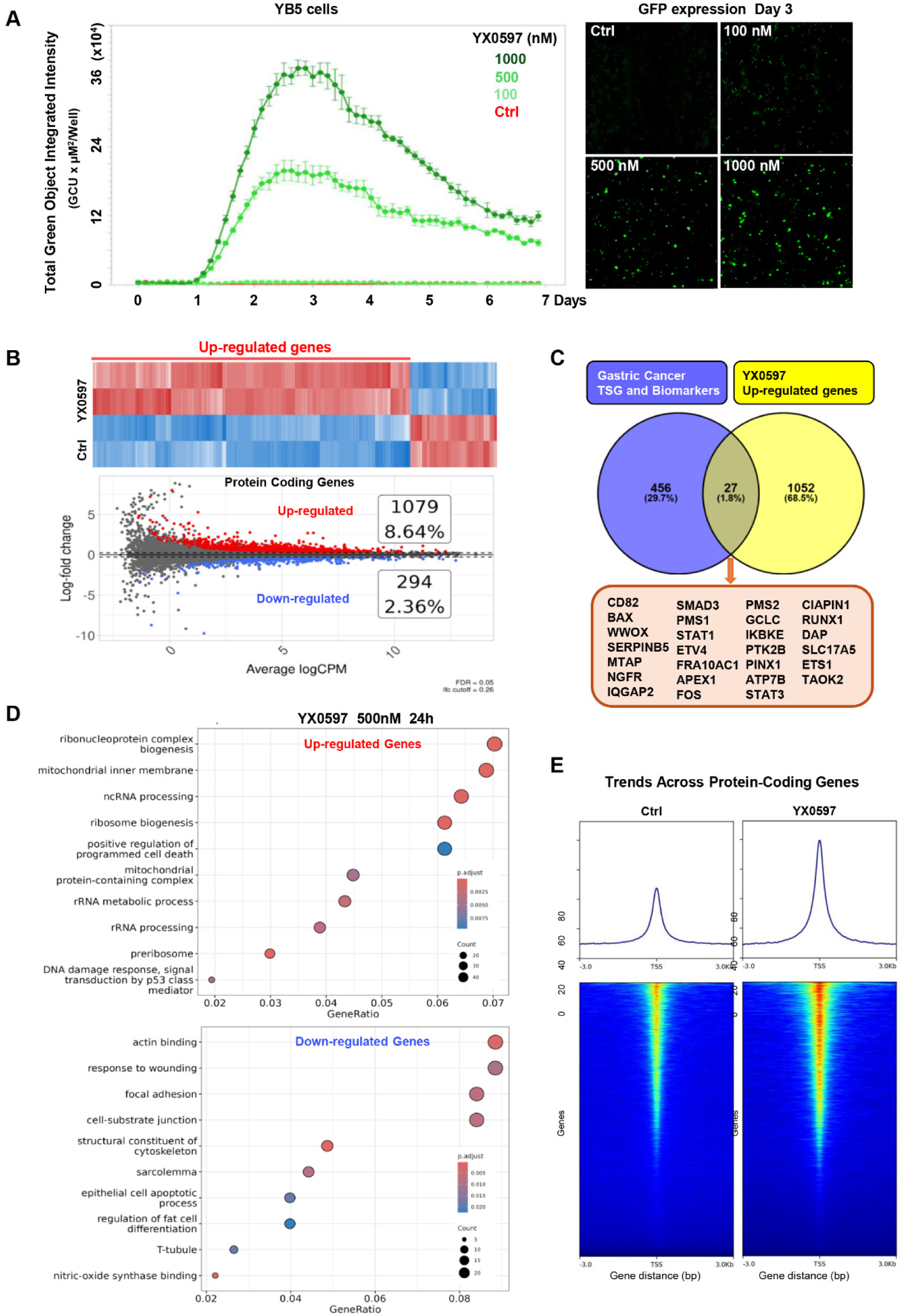
YX0597 reactivates epigenetically silenced genes by increasing chromatin accessibility. **A,** YX0597 dose- and time-dependently activated an epigenetically silenced GFP gene in YB5 cells. Right, GFP fluorescence images after 3 Days treatment. **B,** Bulk RNA-seq analysis of GA0518 cells treated with 500 nM YX0597 for 24 hours. Uppanel, heat map of YX0597 treatment vs. control. **C,** Upregulated Gastric cancer suppressors or biomarkers after YX0597 treatment. **D,** Functional enrichment analysis of the Up-regulated and Down-regulated genes after YX0597 treatment. **E,** ATAC-seq analysis of YX0597 effects on chromatin accessibility of protein-coding genes.

### YAP/TEAD signaling is downregulated by the CDK9 degrader YX0597

As shown in Fig. 3B, RNA-analysis showed ∼ 300 genes downregulated in YX0597-treated cells and many of them were functional enriched in focal adhesion and cell-substrate junction (Fig. 3D). The Hippo signaling pathway is closely linked to cell junctions[16]. YAP and TEAD are proteins in the Hippo signaling pathway that regulate gene expression and cell proliferation. Although normally phosphorylated and degraded, during dyshomeostasis, dephosphorylated YAP1 translocates to the nucleus to transactivate growth-promoting genes, via the transcription factor TEAD[17]. Specifically, YAP/TEAD activity is associated with cancer development, and YAP/TEAD has emerged as an attractive target for cancer therapy [18]. It is worth noticing that multiple YAP/TEAD key downstream effectors were downregulated expression in YX0597 treated cells (Fig. 4A). RT-PCR further validated that YX0597 downregulated the expression YAP/TEAD key downstream effectors Cy61, CTGF, AREG, and Sox9 (Fig. 4B). Public data analysis showed that CDK9 positively correlated with expression of the oncoprotein YAP1, in both EAC and GAC (Fig. 4C), while our single cell analysis showed CDK9^high^ cells highly enriched in YAP^high^ cell group (Fig. 4D). Likewise, IHC staining also supported positive correlation of CDK9 and YAP1, in human gastric adenocarcinoma (Fig. 4E), and that the CDK9 degrader YX0597 dose-dependently decreased YAP1 and TEAD4, at the protein level (Fig. 4F). Immunoprecipitation assays further showed that the transcriptional regulator CDK9 can interact with YAP1 (Fig. 4G) and reciprocally, RNA polymerase II S2 phosphorylation decreased in YAP- knockout cells (Fig. 4H). To further determine whether CDK9 degradation affects YAP1 transcriptional activity, we used a YAP/TEAD luciferase reporter, showing notable downregulation by YX0597 (Fig. 4I). Thus, CDK9 and YAP signaling are closely connected, affecting each another, and downregulated YAP/TEAD signaling contributes to YX0597’s antitumor activity.

**Figure 4.**
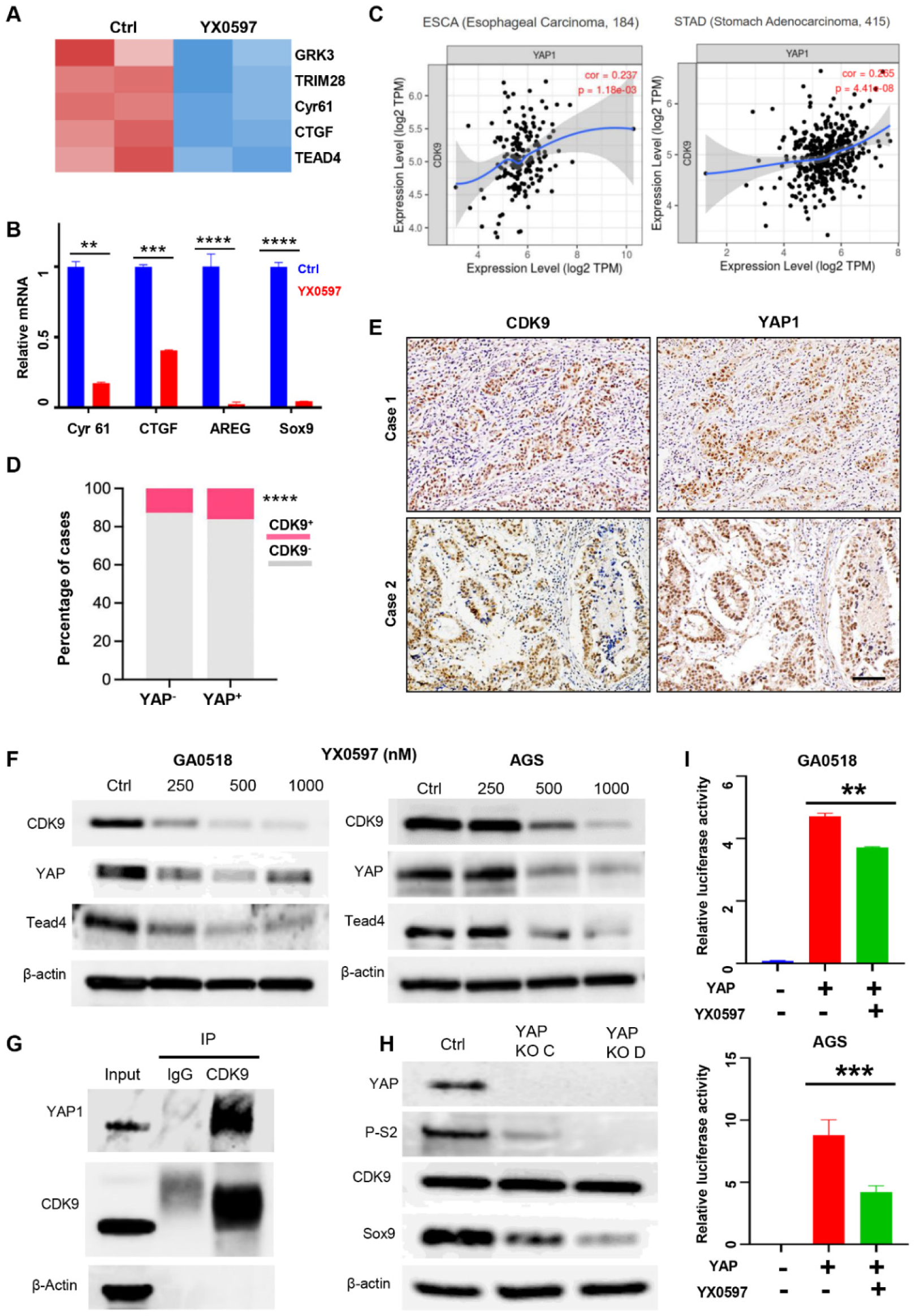
Positive expression correlation of CDK9 and YAP1 expression, and downregulation of YAP-TEAD signaling by the CDK degrader YX0597. **A**, RNA-seq analysis of the effects of YX0597 on YAP-TEAD downstream targets. **B**, RT-PCR analysis of the downstream targets of YAP-TEAD signaling in XY0597-treated cells. **C,** Public database analysis of the expression of CDK9 and YAP1 in GEAC. **D,** Single cell analysis showing CDK9 highly enriched at YAP^high^ cells. **E,** IHC staining of CDK9 and YAP1 using human gastric adenocarcinoma tissues. Scale bar: 100 µm. **F,** Western blotting analysis of the effects of YX0597 on YAP and TEAD4 expression. **G,** Immunoprecipitation (IP) analysis of CDK9 interaction with YAP1. **H,** Effects of YAP1 knockout on phospho-RNA Pol II (Ser2), CDK9, and SOX9. **I,** Effects of YX0597 on YAP-TEAD transcriptional activity. Experiments were performed with three replicates (mean, SD). *\*\*p< 0.01, ***p<0.001, ****p<0.0001*

### Co-targeting CDK9 and YAP/TEAD synergistically inhibits tumor cell growth of YAP^high^ radiation-resistant ESCA cells, both in vitro and in vivo

Given that YAP/TEAD signaling closely correlated with CDK9 activity, we explored whether the novel CDK degrader, YX0597, could sensitize cells to treatment with a YAP/TEAD inhibitor. In particular, cell growth assays showed synergistic growth inhibition by YX0597 in combination with YAP/TEAD inhibitor VT00278 in GA0518 (Fig. 5A), AGS (Fig. 5B) and PDX1004 (Fig. S8) cells. Analogously, cell cycle analysis showed that VT00278 greatly enhanced YX0597 cytotoxicity, as assessed by G2/M phase cells entering apoptosis (Fig. 5C). Moreover, both CDK9 and YAP/TEAD signaling-related effectors markedly decreased in the cotreatment group (Fig. 5D). Therefore, co-targeting CDK9 and YAP/TEAD may represent a new and efficacious means of inhibiting GEAC tumor cell growth.

**Figure 5.**
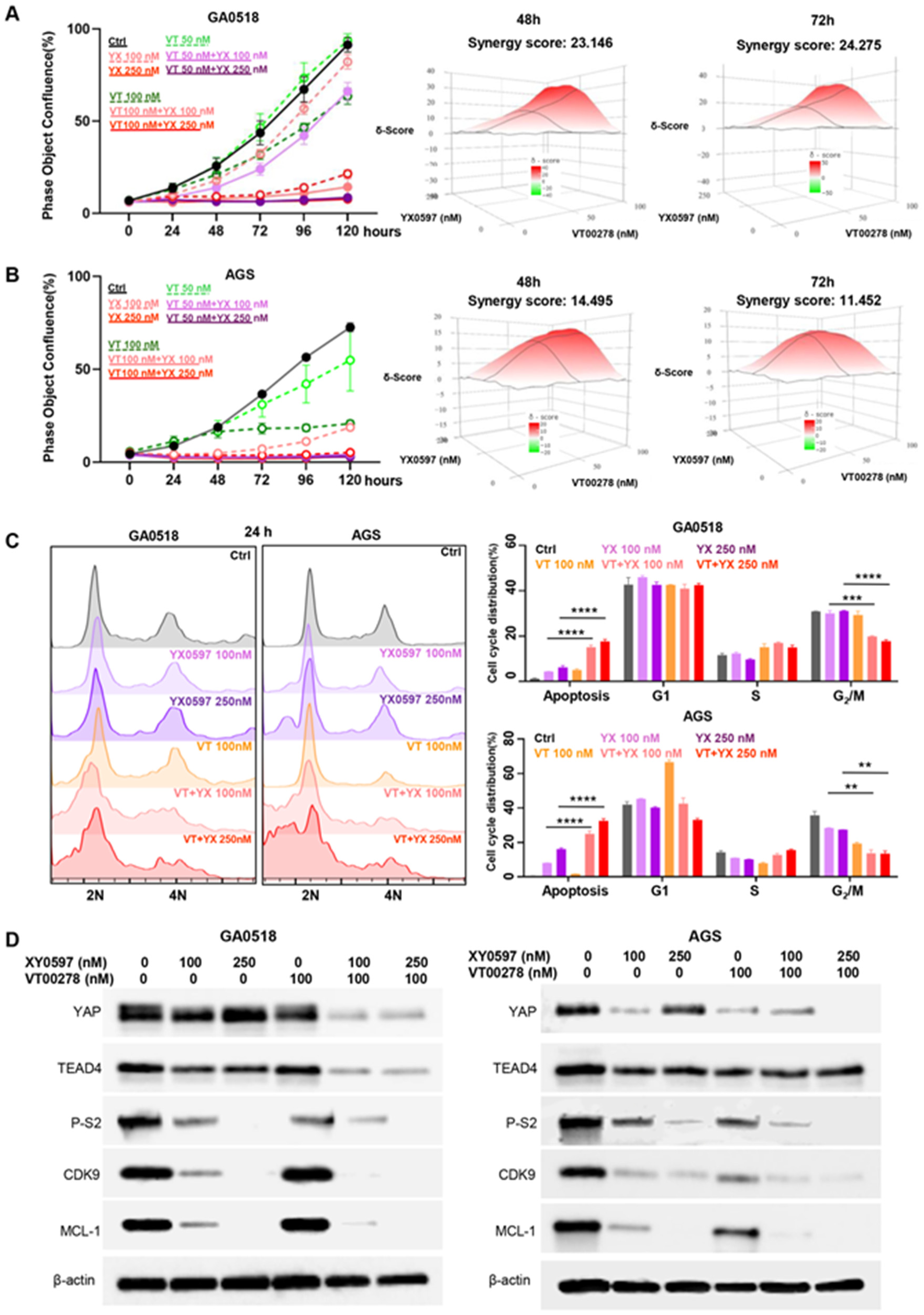
Co-treatment of gastric cancer cells with the CDK9 degrader YX0597 and the YAP/TEAD inhibitor VT00278. **A,** Effects of YX0597, VT00278, or their combination, on GA0518 cell growth (left panel) and synergy scores at 48h (middle) and 72h (right). **B,** Effects of YX0597, VT00278, or their combination, on AGS cell growth (left panel) and synergy scores at 48h (middle) and 72h (right). **C,** Cell cycle analysis of the effects of YX0597, VT00278, or YX0597/VT00278 combination, on GA0518 and AGS cells (left panel). Right panel, quantification by cell cycle phases (\*\**p< 0.01*, \*\*\*\**p<0.0001*). **D,** The effects single and cotreatment of YX0597 and VT00278 on YAP and CDK9 signaling effectors in GA0518 (left) and AGS (right) cells.

Previously, we developed radioresistant Flo-1 XTR cells that showed upregulated YAP1 expression [19], while further confirming a previous finding [20] that, compared to parental (radiosensitive) Flo-1 cells, XTR cells had greater potential to form large tumor spheres (Fig. 6A), an indicator of cancer stemness [21]. Consistent with a role for a CDK9/YAP1 signaling axis, we found that YX0597 powerfully inhibited such tumor sphere generation (Fig. 6A), in addition to inhibiting colony formation (Fig. 6B). Interestingly, YX0597 preferentially decreased CDK9 activity and expression in XTR, vs. parental Flo-1cells (Fig. 6C). Consistent with our current results co-targeting CDK9 and YAP/TEAD in gastric cancer cells (Fig. 5A and 5B), the YAP/TEAD inhibitor VT00278 synergized with YX0597 in Flo-1 esophageal cancer cells (Fig. 6D). Notably, XTR cells were much more sensitive to either single or combination treatment, at low doses (Fig. 6E). Cell cycle analysis further showed that YX0597 induced much more apoptosis in XTR versus parental Flo-1 cells (Fig. 6F). Similar to our data in gastric cancer cells (Fig. 5D), both CDK9 and YAP/TEAD signaling-related effectors were markedly decreased in the cotreatment group (Fig. 6G). Consistent with YX0597 downregulation of SOX9, a downstream target of YAP protein activation [22], in gastric cancer (Fig. 4B), we saw that high SOX9 expression was markedly inhibited, following CDK9 degradation, in XTR cells (Fig. 6G). These data suggest that high YAP activity, in tumor cells, should be more sensitive to YX0597 treatment. Therefore, we injected XTR cells into NSG mice, showing that YX0597 (1 mg/kg) could effectively inhibit xenograft growth, while cotreatment with VT00278 greatly inhibited tumor growth (Fig. 7A-C). Immunofluorescence (IF) staining further showed decreased CDK9, YAP1, KI67, and SOX9, in XTR xenografts, following YX0597 single treatment or combination with VT00278 (Fig. 7D).

**Figure 6.**
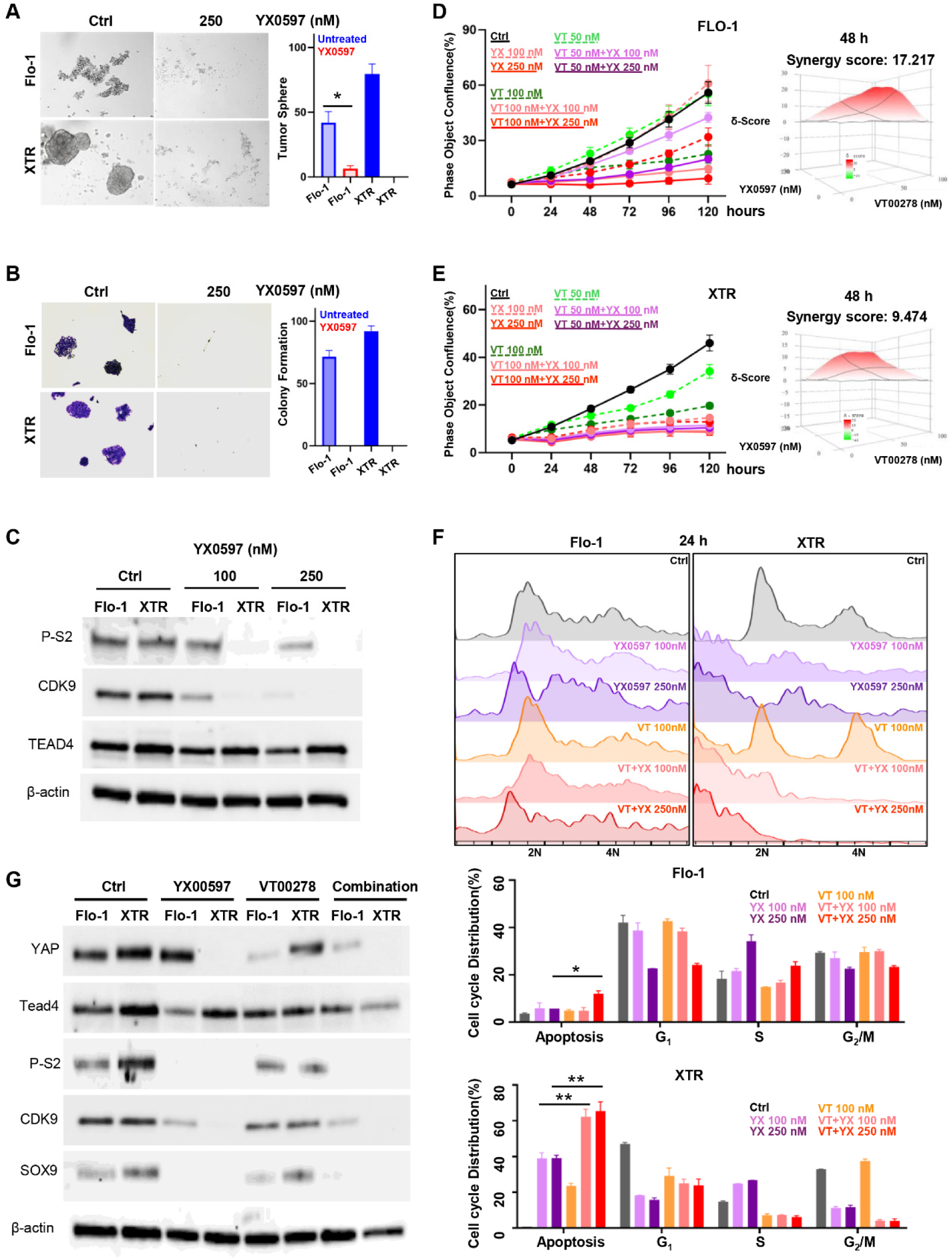
Effects of YX0597, VT00278, or co-treatment on radiation-resistant human esophageal adenocarcinoma cells *in vitro*. Effects of YX0597 on **A,** tumor sphere generation; and **B,** colony formation, of Flo-1 and Flo-XTR cells. *\*p<0.05*. **C,** The effects of YX0597 on phospho-RNA Pol II (Ser2), CDK9, and TEAD4, in Flo-1 and Flo-XTR cells. **D,** Effects of YX0597, VT00278, or co-treatment on Flo-1 cell growth. Right panel, synergy scoring. **E,** Effects of 48h YX0597, VT00278, or co-treatment on Flo-1/XTR cell growth. Right panel, synergy scoring. **F,** Cell cycle effects of 24h YX0597, VT00278, or co-treatment on Flo-1 (left) and Flo-1/XTR (right) cells. Lower panel, quantification. **G,** Effects of YX0597, VT00278, and YX0597/VT00278 treatment on YAP and CDK9 signal mediator expression in Flo-1 and Flo-XTR cells.

**Figure 7.**
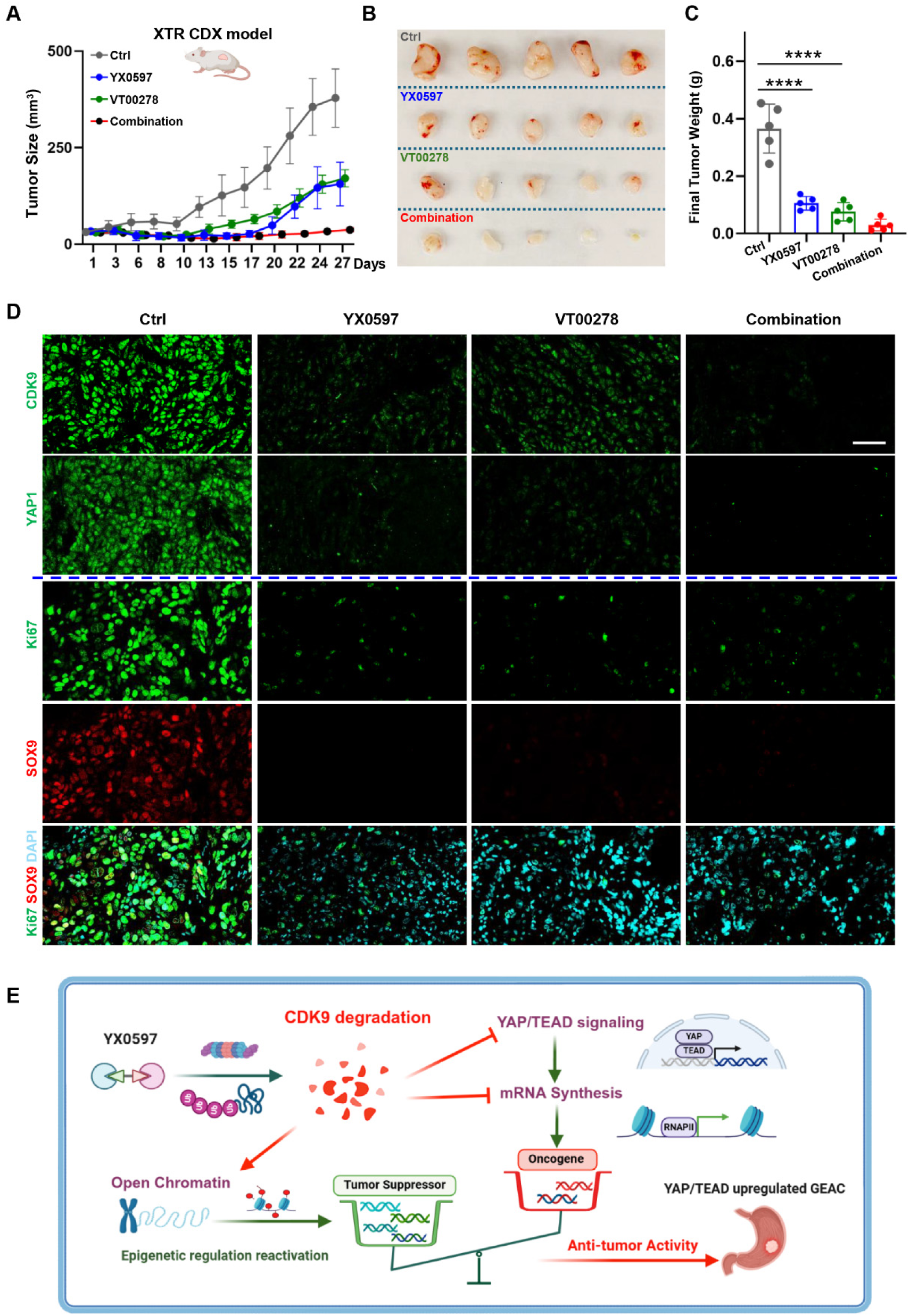
Single or combined treatment effects of YX0597 and VT00278 on Flo-XTR xenograft (CDX) tumor growth. **A,** Effects of YX0597, VT00278, or combination treatment on Flo-XTR xenograft tumor growth. **B,** Images of resected tumors after treatment. **C,** Final tumor weights after treatment. *\*\*\*\*p< 0.001*. **D,** Xenograft tumor immunofluorescence of expression of CDK9, YAP1, Ki67 and SOX9 following control, YX0597, VT00278, and YX0597/VT00278 treatment. Scale bar: 50 nm. **E,** Potential Mechanisms of CDK9 degrader induced cytotoxicity of GEAC tumor cells.

In general, our study showed that novel CDK9 degrader (YX0597) not only remarkably inhibits YAP/TEAD signaling, alone, but also sensitizes GEAC cells to a YAP/TEAD inhibitor. Besides GEAC cells, this combination also demonstrates strong inhibitory effects on YAP^high^ radiation-resistant EAC cells, both *in vitro* and *in vivo*. Mechanistically, YX0597 may downregulate GEAC oncogenes, via decreasing YAP/TEAD signaling and inhibiting RNA pol II phosphorylation, while upregulating key tumor suppressor genes. The loss of balance between these two groups of genes likely contributes to the cytotoxicity induced by YX0597 (Fig. 7E).

## DISCUSSION

Esophageal carcinoma (EAC) and gastric adenocarcinoma (GAC), collectively GEAC, are common and deadly GI tract tumors. While recent research has highlighted pivotal roles for cyclin-dependent kinase-9 (CDK9) in the genesis of several cancers, including proliferation, survival, cell cycle regulation, DNA damage repair, metastasis, and stemness [12], its contribution to GEAC remains unclear. In this study, we found CDK9 highly expressed in both EAC and GAC tumor tissues, in the TCGA database, and IHC further showed much higher CDK9 levels in tumor, compared to stromal cells (Figure 1). To evaluate targeting CDK9 for cancer therapy, we developed a novel CDK9 degrader, YX0597, which strongly suppressed CDK9 and exhibited potent antitumor activity. Mechanistically, YX0597 strongly enhanced chromatin accessibility, activating epigenetically silenced genes and downregulating the oncogenic YAP/TEAD pathway. Moreover, VT00278, a YAP1/TEAD inhibitor, synergized with XY0597 treatment, significantly attenuating radiation-resistant tumor growth and progression, in vivo. Thus, our work provides a potential strategy for targeting CDK9-hyperactivated gastroesophageal cancer.

Recently, transcription-associated CDKs have emerged as important targets in the field of oncology. As a key member of this CDK subfamily, CDK9 plays critical roles in regulating gene expression. In particular, CDK9 inhibition prevents phosphorylation of Serine 2 in the carboxy-terminal domain of RNA Polymerase II (RNA Pol II), preventing transcription elongation of genes critical to the survival and proliferation of cancer cells, including MYC target genes, the lncRNA HOTAIR, PPP1R13L, RBPJ target genes, MCL1, and BIRC5 [12, 23]. Consistent with these reports, using the CDK9 degrader YX0597, we observed rapid and persistent RNA polymerase II P-S2 dephosphorylation, in concert with inhibition of the CDK9 downstream effector MCL1, a Bcl-2 family anti-apoptotic protein. Thus, MCL1 inhibition represents a mechanism of CDK9 degradation-induced apoptosis. Previously, we reported that CDK9 inhibition enables recruitment of the nucleosome remodeler, BRG1, to heterochromatin, facilitating reactivation of a hypermethylated and silenced GFP reporter [9]. In the current study, we used the same YB5 reporter system and detected robust reactivation of silenced GFP, following treatment with two different CDK9 degraders.

Epigenetic gene regulation, characterized by reversibility and susceptibility to external factors, is an efficient way for tumor cells to respond to the environment, and alter gene expression, without changing their primary DNA sequence [24]. In YX0597-treated cells, more than 1000 genes were reactivated and upregulated by increased chromatin accessibility. Interestingly, these upregulated genes included gastric tumor suppressor genes and biomarkers, including *WWOX*, *PINX1*, *IQGAP2*, and *MTAP* (Fig. 3C), in addition to upregulation of transposable elements, a possible means of increasing tumor immunogenicity [25]. RNA polymerase II is well known for transcribing DNA into precursors of messenger RNA (mRNA). Inhibition of RNA polymerase II S2 phosphorylation systematically disturbs numerous genes’ expression patterns in tumor cells, including epigenetic gene reactivation. Consistent with this, our ATAC-seq analysis detected increased genome-wide chromatin accessibility after YX0597 CDK9 degrader treatment. Thus, the tumor cell transcriptome was disturbed after CDK9 degradation. Although some upregulated genes may benefit tumor survival, increased expression of tumor suppressors may contribute to YX0597-induced cytotoxicity (Figure 7E).

Besides upregulated genes, around 300 genes were downregulated in YX0597-treated cells and these genes were functional enriched in focal adhesion and cell-substrate junction which are closely linked to Hippo signaling pathway. We further confirmed that multiple downstream effectors of YAP/TEAD signaling were down-regulated expression. Downregulation of specific transcripts is likely associated with decreased phosphorylation of Pol II Serine 2, interfering with transcriptional elongation, as shown previously in prostate cancer [6]. At the protein level, YAP and TEAD4 were even markedly decreased in YX0597-treated cells. YAP/TEAD signaling is a component of the Hippo organ size control pathway that involves the interaction of Yes-associated protein (YAP), transcriptional coactivator with PDZ-binding motif (TAZ), and transcriptional enhanced associate domain (TEAD) transcription factors. This interaction plays a key role in tumorigenesis, and YAP overexpression is associated with cancer stemness and tumor resistance [22]; thus, pharmacological regulation of YAP/TEAD signaling has potential clinical application in cancer prevention, treatment, and regenerative medicine. We observed the expression of CDK9 and YAP expressions is positively correlated in GEAC patient tumors (Fig 4A-E). Moreover, it has been reported that YAP can recruit a CDK9 complex to enhance expression of YAP-regulated genes [26]. Specifically, YAP/TAZ contacts Pol2 through the Mediator complex (MED1), promoting CDK9-mediated phosphorylation of the Pol2 C-terminal tail, favoring transcriptional pause-release [27]. Consistent with this, we detected CDK9 binding to YAP1 in gastric cancer cells, while interestingly, CDK9 activity could reciprocally be affected in either YAP-KO or VT00278 (YAP inhibitor)-treated cells. Furthermore, our YAP/TEAD luciferase reporter assay supported a close linkage of CDK9 activity to YAP-driven transcriptional activity. Based on this relationship between YAP and CDK9, as expected, the CDK9 degrader YX0597 enhanced YAP inhibition by the YAP/TEAD inhibitor VT00278, with the two drugs demonstrating significant synergy. Interestingly, we also detected that SOX9, a downstream target of YAP, was inhibited by either the CDK9 degrader or YAP/TEAD inhibitor. SOX9 is a transcription factor that has been implicated in both initiation and progression of multiple solid tumors. Together, our work suggests a strong connection between CDK9 and YAP1, and the inhibition of YAP/TEAD signaling represents one mechanism contributing to the cytotoxicity of the CDK9 degrader YX0597.

Another significant finding in this study was the ability of the CDK9 degrader to sensitize radiation-resistant cells. Radiation-resistant cancer is a major obstacle to cancer treatment, leading to cancer recurrence, metastasis, and poor prognosis. Research is ongoing to identify new therapies for radiation-resistant cancer. Previously, we generated radiation-resistant Flo-XTR cells from human esophageal adenocarcinoma FLO-1 cells and further demonstrated highly activated YAP signaling in these cells [20]. Compared to parental Flo-1 cells, Flo-XTR cells were more sensitive to treatment with either YX0597 (CDK9 degrader) or VT0078 (YAP/TEAD inhibitor). Moreover, cotreatment with both drugs demonstrated strong inhibitory effects on Flo-XTR cancer stem-like cells, both *in vitro* and *in vivo*. Thus, co-targeting CDK9 and YAP/TEAD may be a novel treatment strategy for radiation-resistant cancer.

Several CDK9 inhibitors have recently been designed through molecular modeling techniques, showing excellent antitumoral activity *in vitro*, and some CDK9 inhibitors, such as Enitociclib and Dinaciclib, have shown promising results in clinical trials of hematological malignancies [28]. Traditional drug discovery methods repeatedly revolve around developing small molecules designed to inhibit the enzymatic activity of a target protein. While these approaches have yielded prominent successes with different kinase inhibitors, they may not fully ablate the manifold functions of a target protein. Toward that objective, Proteolysis-Targeting Chimeras (PROTACs) represent a significant shift in drug discovery and a promising approach for targeted cancer therapy. The PROTAC recruits an E3 ligase to the protein of interest, triggering its ubiquitination and degradation by the ubiquitin-proteasome system (UPS) [13]. PROTACs have several advantages over traditional anti-cancer therapies, such as the potential to target undruggable proteins, the ability to overcome drug resistance, a prolonged action time, activity in smaller doses, a good safety profile, and more specific selectivity [13, 29]. PROTACs are currently being tested in clinical trials for many types of cancer, including breast and prostate cancer. However, PROTACs can cause on-target toxicities if the protein of interest is not tumor-specific. To address this, researchers are developing PROTACs that recruit E3 ligases enriched in tumors [30]. It has been reported that CDK9 drives vital oncogenic pathways, and it is frequently overexpressed across various types of cancer (Fig.S1A). Here, we first showed that CDK9 is highly expressed in GEAC cancer cells and then synthesized five CDK9 degraders, showing that YX0597, a novel CDK9 degrader designed based on the published CDK9 inhibitor AZD4573 [8], showed potent dose-dependent growth inhibition of multiple GEAC tumor cell lines, and xenograft tumors, at low-doses, *in vivo*. Consequently, we believe YX0597 has the potential to be translated into clinical practice in the future.

In this study, we show that CDK9 is a viable anticancer target in GEAC, Moreover, we demonstrate that a novel CDK9 degrader, YX0597, has high potential clinical application for GEAC treatment, especially in tumors with high YAP signaling activity, such as radiation-resistant cancer. Finally, we reveal crosstalk of CDK9 and YAP/TEAD signaling, and that co-targeting these two mediators could be a novel treatment strategy for the treatment of GEAC.

## METHODS

### Antibodies and chemical reagents

CDK9 antibody (C12F7, #2316), MCL1 antibody (D35A5, #5453), β-Actin antibody (13E5, #93473), Cleaved Caspase-3 antibody (Asp175, #9661), YAP antibody (# 4912; D8H1X #14074) were purchased from Cell Signaling Technology. RNA pol II CTD phospho Ser2 antibody (Active motif, #39563) were supplied by Dr. ISSA group at Coriell. Tead4 antibody (ab155244) were purchased from Abcam. Sox9 antibody(#AB5535) was purchased from Sigma-Aldrich. VT00278 was supplied by Vivace Therapeutics. CDK9 degraders were synthesized by the team of Dr. Jia Zhou (University of Texas Medical Branch).

### Cell lines and cell culture

GA0518 cells were developed from the patient of gastric adenocarcinoma with peritoneal carcinomatosis[15]. AGS cells were purchased from ATCC. The human esophageal adenocarcinoma cell lines OE33 and SKGT4 cells were provided by Dr. Chen group at Coriell. The radiation-resistant (XTR) esophageal adenocarcinoma cell lines Flo-1 XTR and its parental cell line Flo-1 have been described previously [20]. YB5 cells were developed by Dr. Issa team [31]. The cells were cultured in RPMI1640 (Invitrogen, for GA0518 cells) or DMEM (Invitrogen) supplemented with 10% fetal bovine serum (FBS) (Invitrogen), 100 U/ml penicillin, and 100 mg/ml streptomycin (Invitrogen), and maintained at 37°C in a humidified incubator of 5% CO_2_.

### Cell proliferation and Synergy analysis

Live-Cell proliferation analysis was measured using the Incucyte® cell count proliferation assay(Sartorius). Briefly, cells were seeded into 96-well plates. On the next day, culture medium was replaced. Cells were treated and put into the Incucyte system for taking and analysis phase-contrast images of cells. Three independent experiments were performed, each in triplicates. The Synergy effects of drug co-treatment were evaluated by SynergyFinder Plus [32].

### Protein extraction and Western blotting

Cells were rinsed twice with cold PBS and then lysed on ice for 20 minutes in RIPA buffer (Cell Signaling Technology, #9806). After centrifugation at 13,000 ×g for 10 minutes, the protein concentration was measured by Pierce™ BCA Protein Assay Kits (Thermo Scientific, 23225). 50 µg protein/lane proteins were separated by SDS-PAGE by SDS-PAGE, transferred to polyvinylidene difluoride membrane (Bio-Rad), blocked in 5% nonfat milk, and then blotted with the indicated antibodies.

### RNA isolation and real-time PCR

Total RNAs were isolated and purified using the RNeasy Mini Kit (Qiagen, 74106) and converted to cDNA using Luna® Universal One-Step RT-qPCR Kit (New England Biolabs, E3005). mRNA expression was measured using a real-time PCR detection system (Applied Biosystems, QuantStudio 6 Flex) in 96-well optical plates using SsoAdvanced Universal SYBR Green supermix (Bio-Rad,1725275). GAPDH/Gapdh was used as a control.

### Immunohistochemical staining

GAC TMA containing 421 cases of tumor tissues and 408 cases of normal adjacent tissues (NATs) were used for immunohistochemistry (IHC) staining. Briefly, deparaffinized sections were pretreated to retrieve antigens with a Tris-based Antigen Unmasking Solution (Vector Laboratories, Burlingame, CA) before blocking with 10% normal serum and then were incubated with primary antibodies against CDk9 or YAP1 overnight at 4 °C. Immunohistochemical staining was performed using the streptavidin–peroxidase reaction kit with DAB as a chromogen (ABC kit, Vector Laboratories) as previously described[33].

### Analysis of cell cycle and apoptosis via flow cytometry

Cell-cycle analysis was performed as described previously [34]. In brief, cells were fixed with 70% cold ethanol at 20 for at least 1.5 hours and stained with PBS containing 50 mg/mL propidium iodide (PI) and 30 mg/ml RNase A, and the samples analyzed for DNA content and apoptotic cells (sub-G0/G1 fraction) via a flow cytometer (FACSymphony^TM^ A3; Becton Dickinson).

### Colony formation assays

Five hundred cells were seeded into 12-well plates and incubated for 7 days after the appropriate treatment. The colonies were stained with crystal violet (0.5% w/v) and images taken using AMG EVOS Microscope.

### Immunofluorescence microscopy

After fixing the cells using in 4% paraformaldehyde or completing the antigen retrieval of FFPE slides, indirect immunofluorescence staining was done as previously described [35]. Nuclei were revealed by DAPI (1µg/ml) staining. Fluorescence images were observed and collected under Olympus FV3000 Scanning Confocal Microscope.

### Tumor sphere formation assay

Single cells (800/well) were seeded in triplicate onto a 48-well ultra-low attachment plate (Corning) in serum-free DMEM/F-12 supplemented with 10 ng/mL epidermal growth factor, 5 mg/mL insulin, 0.5 mg/mL hydrocortisonum, and bovine pituitary extract (Invitrogen). After 10 to 14 days of culture, the number of tumor spheres formed (diameter >100 mm) was counted under microscope.

### RNA-seq

RNA from experiments was isolated using RNeasy Mini Plus Kit Qiagen, 74106) and sequenced by Novogene. The data was analyzed by the bioinformatic team of Coriell Institute.

### ATAC-seq

ATAC-seq was performed as described [9, 36]. The samples were sequenced by Novogene and analyzed by the bioinformatic team of Coriell Institute.

### Luciferase reporter assays

Transient cotransfection of the plasmids of Flag-YAP, Gal4-TEAD, UAS-luciferase reporter and Renilla vector into GA0518 and AGS cells and the luciferase expression was measured as described previously [37].

### Co-Immunoprecipitation

Endogenous Co-IP experiment was performed in GA0518 cells using CDK9 (Cell Signaling Technology, #2316) and were done as previously described [9].

### *In vivo* xenograft mouse model

Immune-deficient NOD/SCID mice (6-week-old) were purchased from Jackson Laboratories and raised at Cooper Health Care Animal Room. Flo-XTR cells (3×10^6^ cells in 0.1 ml) were suspended in 100 µL Matrigel matrix (Corning, 354234) diluted with PBS at a 1:1 ratio and subcutaneously injected into the backs of the NOD/SCID mice(Male and Female at 1:1 ratio). After the tumors were palpable (tumors reached the approximate size of 5 mm in diameter) mice were randomly divided (5 mice/group) into four groups(Ctrl, YX0597 (intraperitoneal,1 mg/Kg),VT00278 (oral administration,1 mg/Kg), and the Combination of YX0597 and VT00278) for 4 weeks treatment(given every 3 days) as indicated in the figure legends. Tumor volume was calculated by the formula: tumor volume (mm^3^) = 0.5× [length (mm)] × [width (mm)]^2^.

### Patient and public involvement

Patients or the public WERE NOT involved in the design, or conduct, or reporting, or dissemination plans of our research.

### Ethics Statement

All animal experiments were approved by the Institute’s Institutional Animal Care and Use Committee (IACUC). For GAC TMA sections, all patients provided written informed consent, and the study was approved by the ethics committee at the China Medical University.

### Statistical analysis

Data are presented as mean ± SD. Statistical analysis was performed using GraphPad Prism version 10.0 (GraphPad Software Inc., San Diego, CA). The statistical difference between two comparing groups was compared using a Student’s t-test. One-way ANOVA followed by Dunnett’s multiple comparison test was used to analyze the statistical significance of 3 or more groups. P value <0.05 is considered statistically significant.

## Data Availability Statement

Coriell institute for medical research is committed to data sharing. All scientific data including raw/measured and derived data will be preserved at the institute computing facility. Relevant data will be shared upon publication for the purposes of reproducibility and reusability. The metadata and other relevant data will be deposited to a public repository such as the Gene Expression Omnibus (GEO) database, which provides free access to the scientific community.

## Authors’ Disclosures

The authors declare no potential conflicts of interest.

## Authors’ Contributions

**Yan-ting Yann Zhang**: Investigation, Methodology, Formal analysis, Data curation, Writing – original draft, review & editing. **Mikel Ghelfi**: Investigation, Methodology, Formal analysis. **Yu Xue**: Methodology, Resources. **Haiying Chen**: Methodology, Resources. **Gennaro Calendo**: Methodology, Formal analysis, Data curation. **Generosa Grana**: Resources, Project administration. **Yuan Li**: Investigation, Formal analysis. **Dipti Athavale**: Investigation, Formal analysis. **Curt Balch**: Writing – original draft, review & editing, Formal analysis. **Xiaodan Yao**: Investigation, Resources. **Woonbok Chung**: Methodology, Formal analysis. **Zhenning Wang**: Resources, Project administration. **Francis Spitz**: Resources, Project administration. **Tracy Tang**: Resources, Methodology. **Vladimir Khazak**: Resources, Methodology. **Jia Zhou**: Methodology, Supervision, Resources. **Jean-Pierre Issa**: Supervision, Resources, Project administration, Formal analysis. **Shumei Song**: Supervision, Resources, Project administration, Funding acquisition, Formal analysis, Data curation, Writing – original draft, review & editing.

## Acknowledgements

This work was supported by grants from the NIH (R01 CA269685) and DOD (CA210457 and CA230323).

## Supplementary Figures

**Figure S1.**
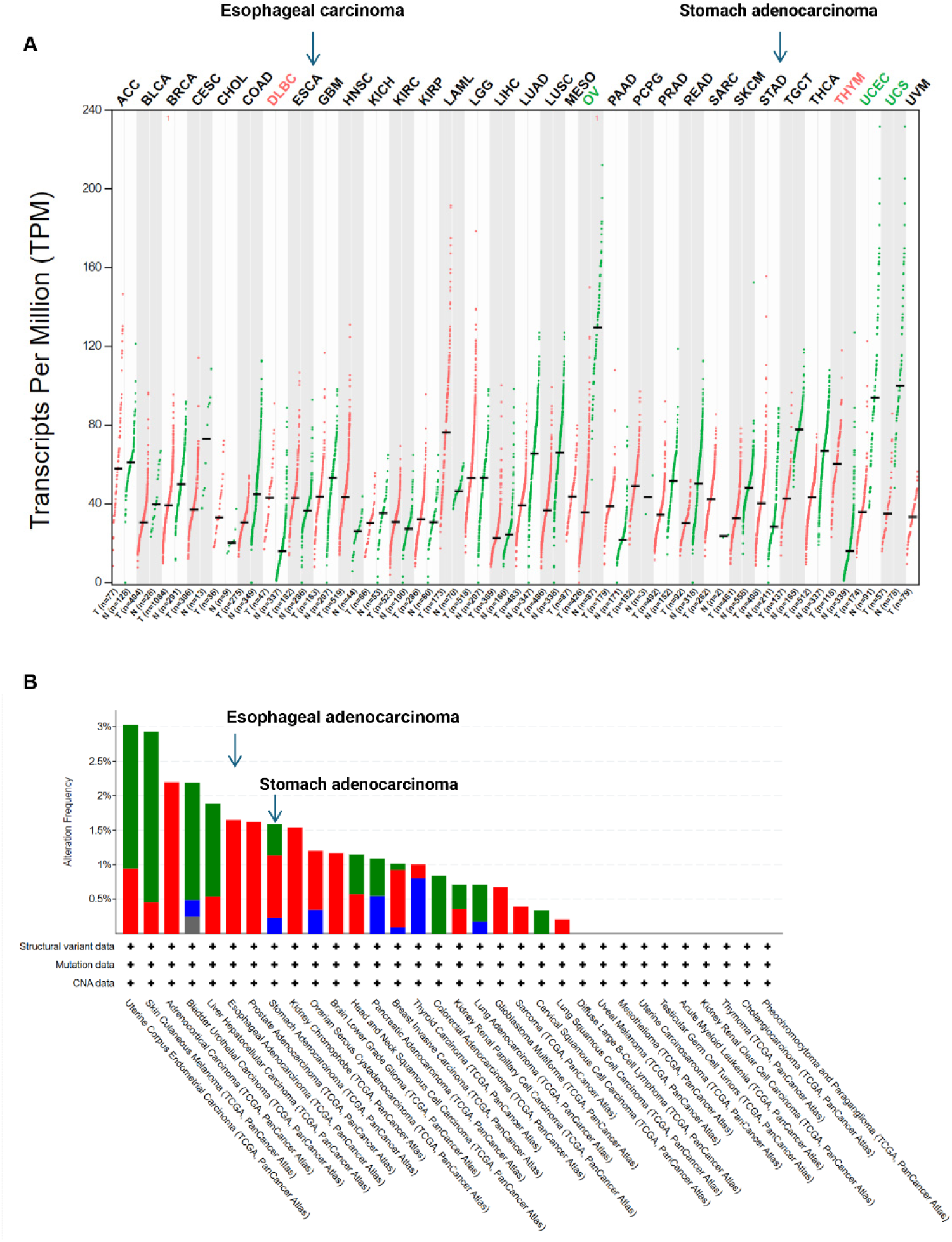
Public datasets of the CDK9 expression across all tumor samples and paired normal tissues (**A**) and genetic alterations profile(**B**).

**Figure S2.**
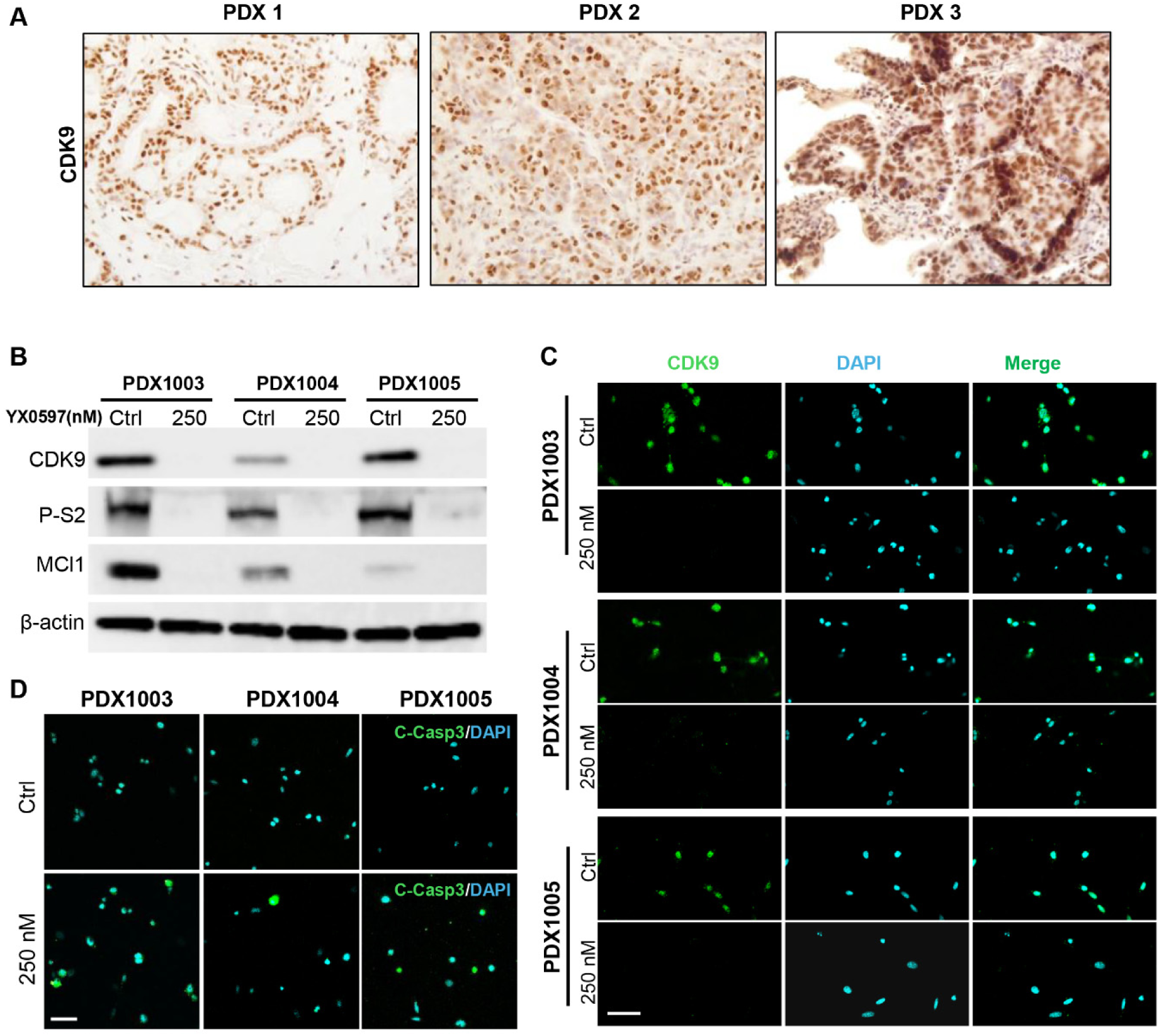
The distribution of CDK9 in PDX tissue and the effects of YX0597 on CDK9 signaling of PDX cells. **A,** Immunohistochemistry of CDK9 expression in PDX tissues. **B,** Western blotting analysis the effects of YX0597 on CDK9 signaling. **C,** Immunofluorescence staining checking the effects of YX0597 on CDK9 degradation. **D,** Immunofluorescence staining checking the effects of YX0597 on Caspase 3 cleavage(activation).

**Figure S3.**
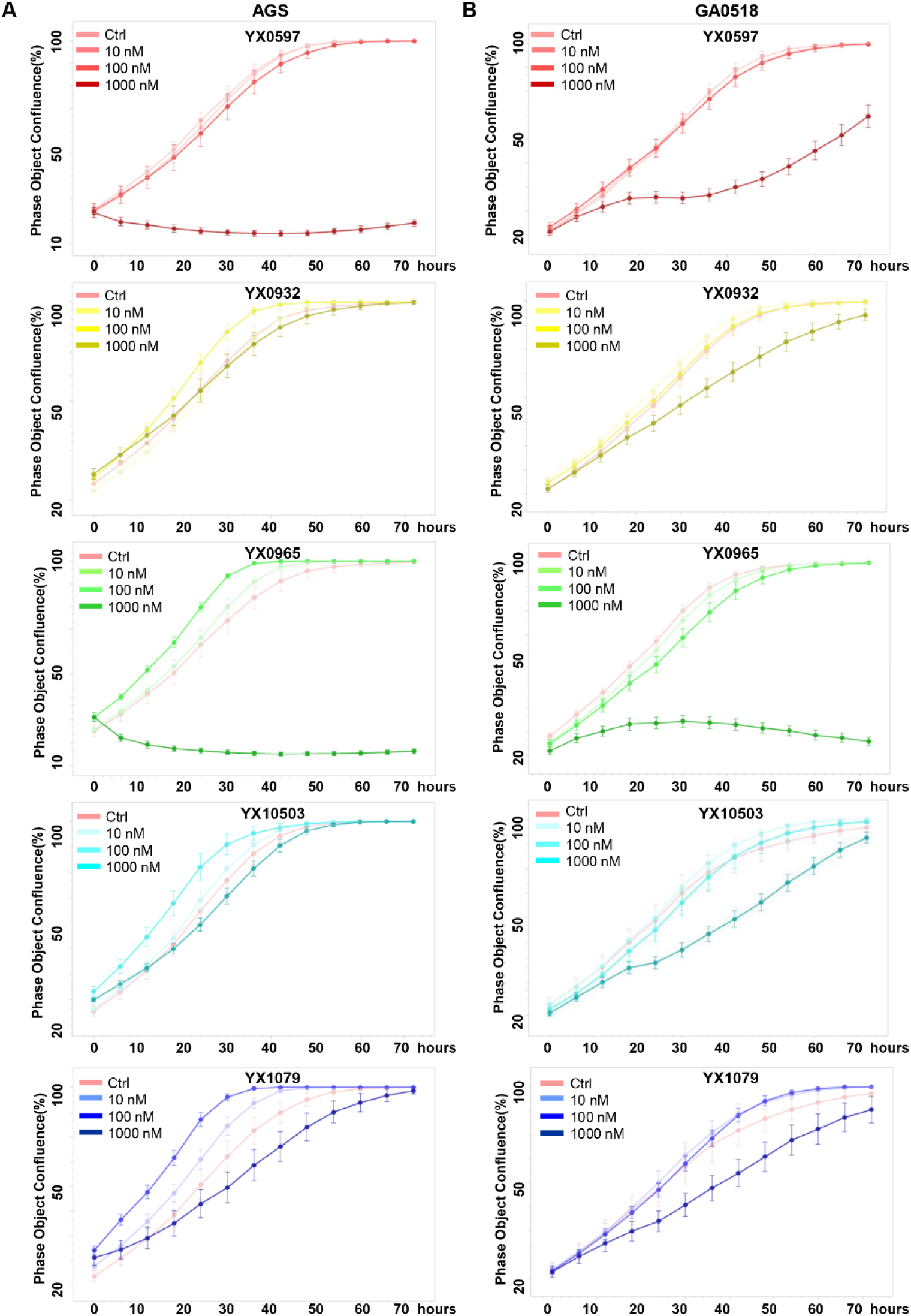
Incucyte live-cell analysis the growth inhibition effects of five novel CDK9 degraders in GA0518 and AGS gastric cancer cells.

**Figure S4.**
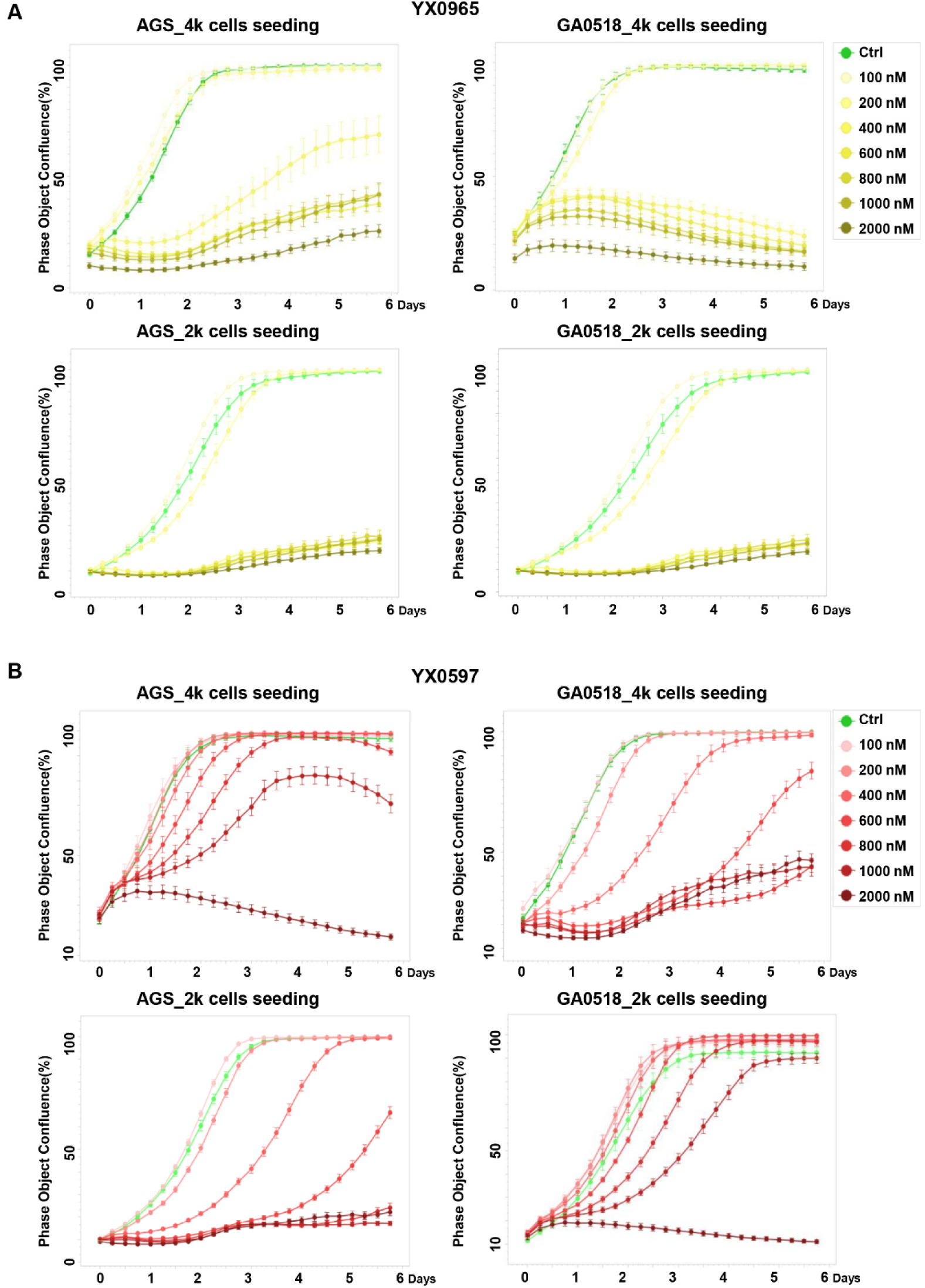
Incucyte live-cell analysis comparison of the growth inhibition effects of CDK9 degrader YX0965 and YX0597 in GA0518 and AGS gastric cancer cells.

**Figure S5.**
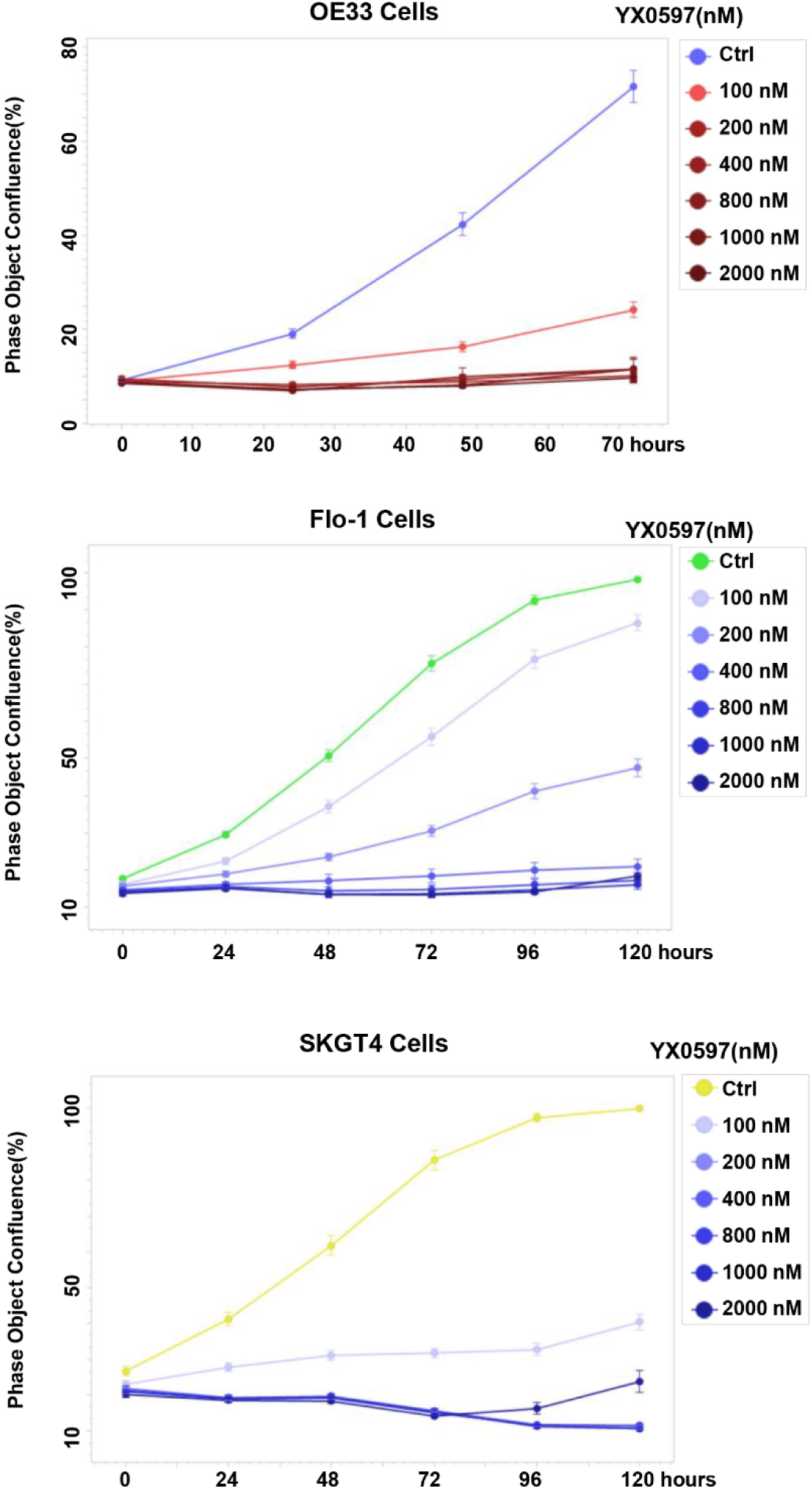
Incucyte live-cell analysis the growth inhibition effects of CDK9 degrader YX0597 in esophageal cancer cell OE33, Flo-1 and SKGT4.

**Figure S6.**
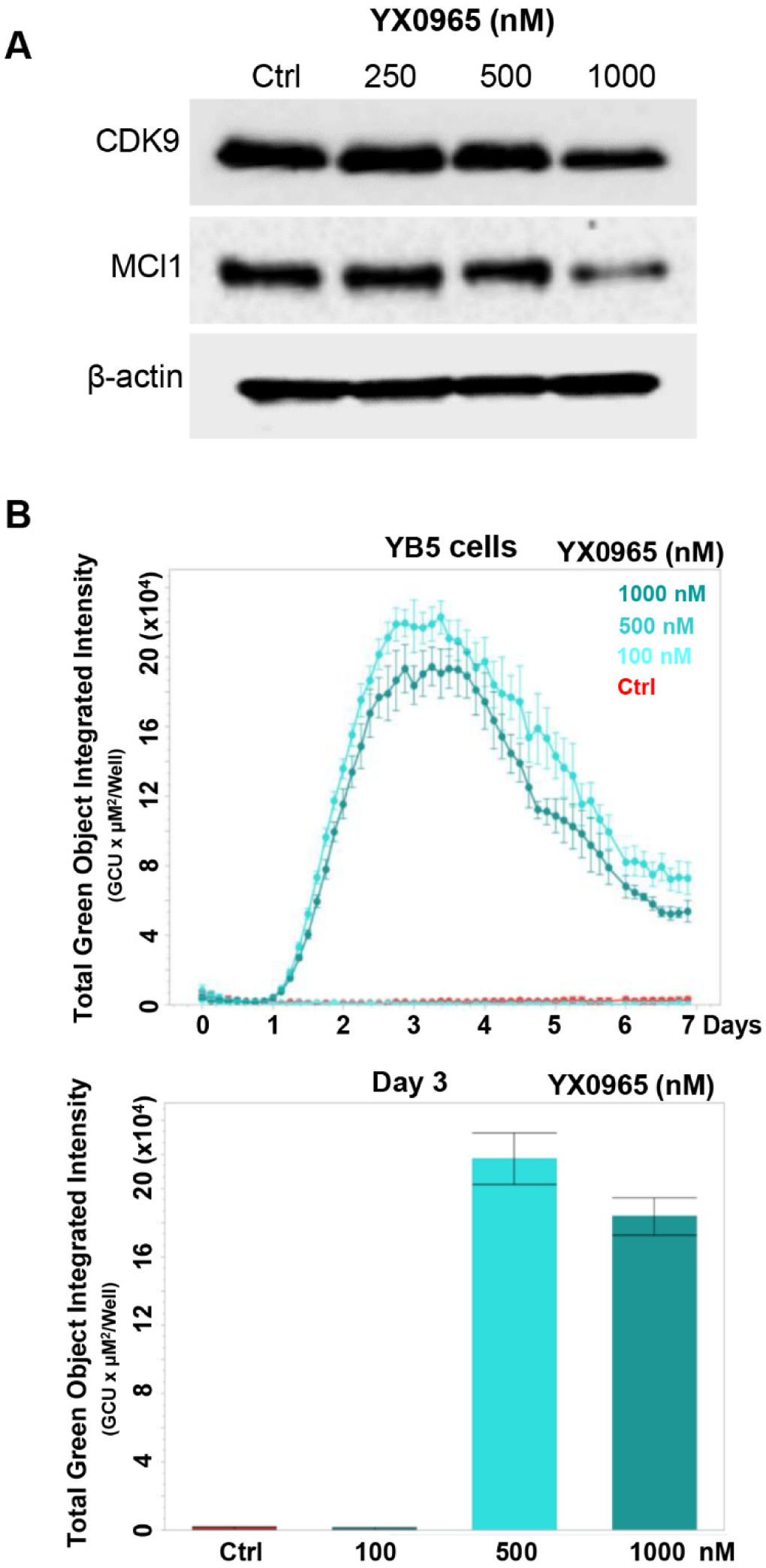
Effects of YX0965 on CDK9 degradation(**A**) and the activation of epigenetically silenced GFP gene in YB5 cells(**B**).

**Figure S7.**
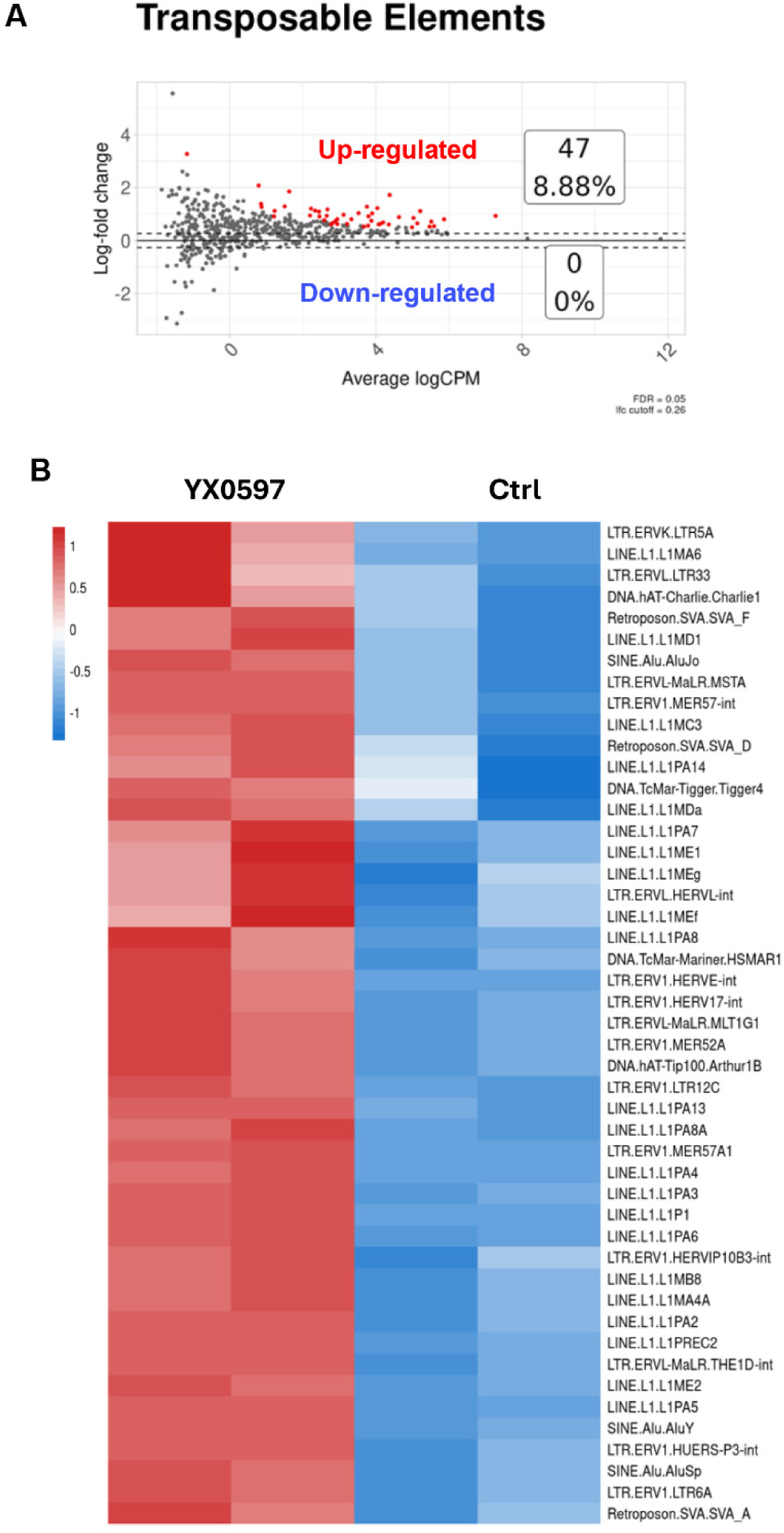
RNA-seq analysis of the transposable elements’ expression in YX0597- treated GA0518 cells.

**Figure S8.**
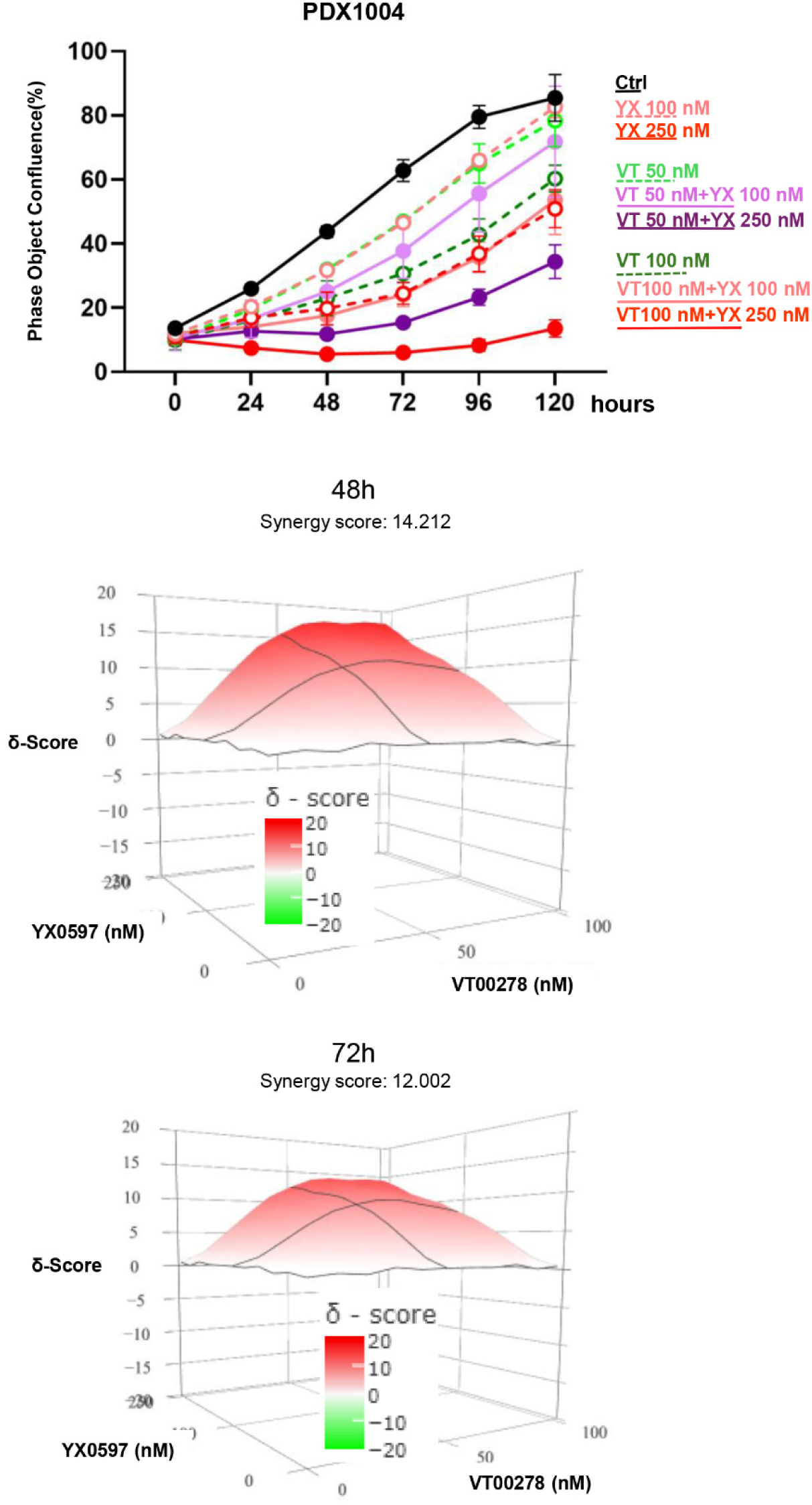
The combination effects of YX0597 and VT00278 on PDX1004 cell growth.

## Notes

### Competing Interest Statement

The authors have declared no competing interest.

## References

1. Global Burden of Disease Cancer, C., et al., *Global, Regional, and National Cancer Incidence, Mortality, Years of Life Lost, Years Lived With Disability, and Disability-Adjusted Life-Years for 29 Cancer Groups*, *1990 to 2016: A Systematic Analysis for the Global Burden of Disease Study*. JAMA Oncol, 2018. 4(11): p. 1553–1568.

2. Thrift, A.P., T.N. Wenker, and H.B. El-Serag, Global burden of gastric cancer: epidemiological trends, risk factors, screening and prevention. Nat Rev Clin Oncol, 2023. 20(5): p. 338–349.

3. Zhang, Y., Epidemiology of esophageal cancer. World J Gastroenterol, 2013. 19(34): p. 5598–606.

4. Anshabo, A.T., et al., CDK9: A Comprehensive Review of Its Biology, and Its Role as a Potential Target for Anti-Cancer Agents. Front Oncol, 2021. 11: p. 678559.

5. Mo, C., et al., CDK9 inhibitors for the treatment of solid tumors. Biochem Pharmacol, 2024. 229: p. 116470.

6. Rahman, R., et al., CDK9 inhibition inhibits multiple oncogenic transcriptional and epigenetic pathways in prostate cancer. Br J Cancer, 2024. 131(6): p. 1092–1105.

7. Ettl, T., D. Schulz, and R.J. Bauer, The Renaissance of Cyclin Dependent Kinase Inhibitors. Cancers (Basel), 2022. 14(2).

8. Cidado, J., et al., AZD4573 Is a Highly Selective CDK9 Inhibitor That Suppresses MCL-1 and Induces Apoptosis in Hematologic Cancer Cells. Clin Cancer Res, 2020. 26(4): p. 922–934.

9. Zhang, H., et al., Targeting CDK9 Reactivates Epigenetically Silenced Genes in Cancer. Cell, 2018. 175(5): p. 1244–1258 e26.

10. Becker, P.B. and J.L. Workman, Nucleosome remodeling and epigenetics. Cold Spring Harb Perspect Biol, 2013. 5(9).

11. Thieme, E., et al., CDK9 inhibition induces epigenetic reprogramming revealing strategies to circumvent resistance in lymphoma. Mol Cancer, 2023. 22(1): p. 64.

12. Mandal, R., S. Becker, and K. Strebhardt, Targeting CDK9 for Anti-Cancer Therapeutics. Cancers (Basel), 2021. 13(9).

13. Bekes, M., D.R. Langley, and C.M. Crews, PROTAC targeted protein degraders: the past is prologue. Nat Rev Drug Discov, 2022. 21(3): p. 181–200.

14. Xiao, L., et al., Targeting CDK9 with selective inhibitors or degraders in tumor therapy: an overview of recent developments. Cancer Biol Ther, 2023. 24(1): p. 2219470.

15. Song, S., et al., Patient-derived cell lines and orthotopic mouse model of peritoneal carcinomatosis recapitulate molecular and phenotypic features of human gastric adenocarcinoma. J Exp Clin Cancer Res, 2021. 40(1): p. 207.

16. Karaman, R. and G. Halder, Cell Junctions in Hippo Signaling. Cold Spring Harb Perspect Biol, 2018. 10(5).

17. Yin, F., et al., Hippo-YAP signaling in digestive system tumors. Am J Cancer Res, 2021. 11(6): p. 2495–2507.

18. Dey, A., X. Varelas, and K.L. Guan, Targeting the Hippo pathway in cancer, fibrosis, wound healing and regenerative medicine. Nat Rev Drug Discov, 2020. 19(7): p. 480–494.

19. Li, F., et al., YAP1-Mediated CDK6 Activation Confers Radiation Resistance in Esophageal Cancer - Rationale for the Combination of YAP1 and CDK4/6 Inhibitors in Esophageal Cancer. Clin Cancer Res, 2019. 25(7): p. 2264–2277.

20. Song, S., et al., A Novel YAP1 Inhibitor Targets CSC-Enriched Radiation-Resistant Cells and Exerts Strong Antitumor Activity in Esophageal Adenocarcinoma. Mol Cancer Ther, 2018. 17(2): p. 443–454.

21. Chen, S.F., et al., Nonadhesive culture system as a model of rapid sphere formation with cancer stem cell properties. PLoS One, 2012. 7(2): p. e31864.

22. Wang, L., et al., Unbalanced YAP-SOX9 circuit drives stemness and malignant progression in esophageal squamous cell carcinoma. Oncogene, 2019. 38(12): p. 2042–2055.

23. Ranjan, A., et al., Targeting CDK9 for the Treatment of Glioblastoma. Cancers (Basel), 2021. 13(12).

24. Herceg, Z. and T. Vaissiere, Epigenetic mechanisms and cancer: an interface between the environment and the genome. Epigenetics, 2011. 6(7): p. 804–19.

25. Wang, X., Q. Zhang, and X. Cao, Reversing epigenetic repression of transposable elements for improving tumor immunogenicity. Cancer Commun (Lond), 2022. 42(3): p. 266–268.

26. Battilana, G., F. Zanconato, and S. Piccolo, Mechanisms of YAP/TAZ transcriptional control. Cell Stress, 2021. 5(11): p. 167–172.

27. Galli, G.G., et al., YAP Drives Growth by Controlling Transcriptional Pause Release from Dynamic Enhancers. Mol Cell, 2015. 60(2): p. 328–37.

28. Morillo, D., G. Vega, and V. Moreno, CDK9 INHIBITORS: a promising combination partner in the treatment of hematological malignancies. Oncotarget, 2023. 14: p. 749–752.

29. Neklesa, T.K., J.D. Winkler, and C.M. Crews, Targeted protein degradation by PROTACs. Pharmacol Ther, 2017. 174: p. 138–144.

30. Khan, S., et al., PROteolysis TArgeting Chimeras (PROTACs) as emerging anticancer therapeutics. Oncogene, 2020. 39(26): p. 4909–4924.

31. Si, J., et al., Chromatin remodeling is required for gene reactivation after decitabine-mediated DNA hypomethylation. Cancer Res, 2010. 70(17): p. 6968–77.

32. Zheng, S., et al., SynergyFinder Plus: Toward Better Interpretation and Annotation of Drug Combination Screening Datasets. Genomics Proteomics Bioinformatics, 2022. 20(3): p. 587–596.

33. Li, Y., et al., Lymphatic Drainage System and Lymphatic Metastasis of Cancer Cells in the Mouse Esophagus. Dig Dis Sci, 2023. 68(3): p. 803–812.

34. Zhang, Y., et al., Cucurbitacin B induces rapid depletion of the G-actin pool through reactive oxygen species-dependent actin aggregation in melanoma cells. Acta Biochim Biophys Sin (Shanghai), 2011. 43(7): p. 556–67.

35. Zhang, Y., et al., Formation of cofilin-actin rods following cucurbitacin-B-induced actin aggregation depends on Slingshot homolog 1-mediated cofilin hyperactivation. J Cell Biochem, 2013. 114(10): p. 2415–29.

36. Buenrostro, J.D., et al., Transposition of native chromatin for fast and sensitive epigenomic profiling of open chromatin, DNA-binding proteins and nucleosome position. Nat Methods, 2013. 10(12): p. 1213–8.

37. Song, S., et al., Galectin-3 mediates nuclear beta-catenin accumulation and Wnt signaling in human colon cancer cells by regulation of glycogen synthase kinase-3beta activity. Cancer Res, 2009. 69(4): p. 1343–9.

